# Aberrant expression of stress-related Hsrω-n lncRNA contributes to CGG repeat-mediated toxicity in a *Drosophila* model of FXTAS

**DOI:** 10.64898/2026.07.10.737441

**Authors:** Nadeem Ahmed, Mudasir Manzoor Reshi, Anand Kumar Singh, Yulin Jin, Peng Jin, Abrar Qurashi

## Abstract

Fragile X-associated tremor/ataxia syndrome (FXTAS) is an adult-onset neurodegenerative disease associated with carriers of premutation (PM) alleles of the fragile X messenger ribonucleoprotein 1 (FMR1) gene. Expanded CGG repeats in the PM alleles are sufficient to cause cellular stress and toxicity in animal and cellular models of FXTAS. Here, we show that in a *Drosophila* transgenic model of FXTAS, expanded CGG repeats result in the overexpression of heat shock RNA omega-nuclear (*Hsrω-n*), a stress-related long non-coding RNA (lncRNA) that dominantly enhances expanded CGG-induced neurotoxicity, whereas its reduction suppresses CGG-induced neurotoxicity. Furthermore, overexpression of *Hsrω-n* lncRNA concomitantly resulted in a significant enhancement in its association with Hrb87F, the fly ortholog of one of the previously identified CGG repeat RNA-binding proteins, hnRNP A2/B1, and with the imitation switch (ISWI) protein, a chromatin remodeling factor. We extended the findings of *Hsrω-n* to show that the mammalian stress-related lncRNAs *SatIII* and *Neat1* are elevated in cells expressing expanded CGG repeats. Together, these findings highlight the critical roles of stress-related lncRNAs in CGG repeat-mediated toxicity and support a model in which the total steady-state levels of stress-related lncRNAs increase in the presence of expanded CGG repeats as part of the normal regulatory response to combat stress. However, the effects of chronic expanded CGG repeat expression in post-mitotic neurons may result in altered distribution and/or sequestration of specific RNA-binding proteins or chromatin remodeling factors associated with stress-related lncRNAs. This could contribute to the altered cellular homeostasis and genomic instability associated with FXTAS.

**Significance:** Individuals carrying the FMR1 allele with expanded CGG repeats (55–200) in the 5′ untranslated region are predisposed to fragile X–associated tremor/ataxia syndrome (FXTAS), a late-onset neurodegenerative disorder. FXTAS demonstrates incomplete penetrance, suggesting the possible involvement of genetic modifiers in its pathogenesis. We used a transgenic *Drosophila* FXTAS model to conduct genetic, molecular, and biochemical analyses. Our findings revealed that the expression of expanded CGG repeats aberrantly activates stress-related Hsrω-n lncRNA in *Drosophila melanogaster.* Overexpression of hsrω-n lncRNA exacerbates CGG-mediated neurodegeneration, whereas its repression alleviates this effect. Extending these findings to mammalian systems, we observed the upregulation of SATIII and Neat I, functional homologs of Hsrω-n lncRNA, in cells expressing expanded CGG repeats. Collectively, these findings identify stress-related lncRNAs as key mediators of CGG-mediated toxicity and underscore their potential as therapeutic targets for FXTAS.

## Introduction

Fragile X–associated tremor/ataxia syndrome (FXTAS) is a late-onset, neurodegenerative debilitating disorder characterized by cerebellar dysfunction, with the main clinical presentations of intention tremor and cerebellar gait ataxia. Typically, in the general population, CGG repeats in the 5′ UTR of the fragile X messenger ribonucleoprotein 1 (FMR1) gene range between five and 54 repeats (1,2). Individuals with more than 200 CGG repeats (full mutation (FM)) develop fragile X syndrome (FXS), a neurodevelopmental disorder (4). On the other hand, carriers of the premutation (PM) allele (55-200 CGG repeats) develop fragile X-associated tremor/ataxia syndrome (FXTAS) with an estimated prevalence of about 1 in 3,000 men and 1 in 8,000 women (7–9). A persisting conundrum is that not all premutation carriers develop FXTAS; ∼40% of males and 16% of females develop FXTAS in late adulthood (1, 2). The incomplete penetrance of FXTAS has motivated the search for genetic modifiers that may serve as genetic biomarkers for prognosis and potential therapeutic targets (10).

Two mutually non-exclusive molecular mechanisms have been proposed for FXTAS. An RNA gain-of-function mechanism is based on the observation of increased levels of FMR1 mRNA containing expanded CGG repeats in the PM range, along with either no detectable change in the FMR1 encoded protein, FMRP, or slightly reduced FMRP levels in PM carriers (17–19). The absence of FXS, which results from FMRP loss, in FXTAS patients, along with the absence of FXTAS symptoms in older individuals with FXS, suggests a direct role of expanded ribo-CGG (rCGG) in FXTAS pathology. Elevated levels of rCGG sequester specific subsets of RNA-binding proteins (RBPs) and limit their availability for normal cellular functions(16,20–22). Another proposed mechanism is repeat-associated non-AUG (RAN) translation, in which rCGG and/or antisense rGCC repeats are non-canonically translated into multiple toxic homopolymeric peptides such as FMRpolyG, the length of which correlates with the number of CGG repeats (23–26). Such homopolymeric peptides have been detected in intranuclear inclusions of PM transgenic animal models. In differentiated neurons derived from FXTAS-induced pluripotent stem (iPS) cells, FMRpolyG disrupts the architecture of the nuclear lamina because of its interaction with the transmembrane protein lamina-associated polypeptide 2 beta (LAP2β) (27). In both models, neurotoxicity arising from the expanded CGG-repeat RNA is of central importance (15,26–30). Therefore, it is critical to determine the mediators of rCGG neurodegeneration.

Several RNA-binding proteins (RBPs) have been shown to interact with rCGG repeats *in vitro* and *in vivo*, with a subset having the potential to influence CGG repeat-mediated toxicity. RBPs that are sequestered directly or in trans by rCGG repeats include hnRNP A2/B1, Pur α (20,21), DROSHA-DGCR8(22), Sam68(13), several serine/arginine-rich domain (SR) proteins(31), CUGBP1 (20), and RNA helicase Rm62(32). hnRNP A2/B1, a multifunctional heterogeneous nuclear RNA-binding protein involved in mRNA metabolism and transport, is particularly intriguing. hnRNP A2/B1 has been consistently identified within ubiquitin-positive intranuclear inclusions in FXTAS patient brains and model systems (10,14). Furthermore, overexpression of human hnRNP A2/B1 and *Drosophila* homologs of the mammalian hnRNP A2/B1 protein, Hrb87F, or Hrb98DE in neurons can suppress neurodegeneration in transgenic *Drosophila* expressing expanded CGG repeats (20,33).

A particular class of stress-related long noncoding RNAs termed "architectural RNAs" (arcRNA’S) functions as scaffolds for some nuclear bodies (34). These include the *Drosophila Hsrω-n* lncRNA in the formation of omega speckles (ω-speckles) (35), human *SatIII* lncRNAs in the formation of nuclear stress bodies (nSB), and nuclear paraspeckle assembly transcript 1 (*Neat 1*) lncRNA in the formation of paraspeckles (34). Although these nuclear bodies have characteristic compositions and functions, they share certain features (34, 36). First, they share some protein factors, such as prion-like domain (PLD)-containing RNA-binding proteins (37) and switch/sucrose non-fermentable (SWI/SNF) chromatin remodeling factors (34). Second, they control the trafficking and availability of transcription and RNA-processing factors to maintain cellular homeostasis under both normal and stressful conditions. Finally, they serve as molecular hubs for cellular processes commonly affected by neurodegenerative diseases such as Huntington’s disease (HD), amyotrophic lateral sclerosis (ALS), and frontotemporal lobar degeneration with ubiquitinated inclusions (FTLD-TDP) (38).

In this study, we showed that the expression of expanded CGG repeats aberrantly activates Hsrω-n expression in a *Drosophila* transgenic model of FXTAS. Our genetic experiments showed that overexpression of Hsrω-n lncRNA enhanced neurotoxicity associated with CGG repeat expression. Similarly, repression of Hsrω-n lncRNA suppressed CGG repeat neurotoxicity. Consistent with this, our genetic experiments showed that Hrb87F mediates the genetic interaction between CGG repeats and *Hsrω-n*. Biochemical experiments revealed that in the presence of CGG repeats, cellular Hrb87F and chromatin remodeling factor, imitation switch (ISWI), were significantly associated with *Hsrω-n*. Immunocytochemistry showed accumulation of Hrb87F to the chromosomal site of *Hsrω-n* synthesis to assemble ω-speckles. Together, our study proposes a model in which nuclear Hrb87F and possibly ISWI, in the presence of CGG repeats, bind to overexpressed *Hsrω-n* lncRNA molecules, thereby modulating CGG-mediated neurodegeneration. We further showed that *Sat III* and *Neat I* expression was increased in mammalian cells expressing expanded CGG repeats. Because *Sat III* and *Neat I* are functional homologs of *Hsrω-n* lncRNAs, stress-related lncRNAs may therefore play critical roles in CGG-mediated toxicity and could potentially provide therapeutic insights into FXTAS.

## Results

### Expanded CGG repeats upregulate expression of *Hsrω-n* lncRNA in the fly brain

In *Drosophila*, the nucleus retained *Hsrω* lncRNA (Fig. 1A) dynamically responds to normal physiological processes and various forms of stress (39). We previously showed that transgenic flies carrying fragile X PM CGG repeats experienced chronic stress and increased expression and nuclear retention of mRNAs from stress response genes (32). Therefore, we examined the expression levels of *Hsrω-n* lncRNA in transgenic flies expressing expanded CGG repeats (15, 32). The transgenes were driven by a pan-neuronal *Elav-*GAL4 driver using the *Drosophila* GAL4/UAS system. Expression of the strong (*pUAST-CGG_90_-EGFP*) transgene in neurons leads to lethality in flies that do not reach adulthood. However, flies carrying the heterozygous transgene *(pUAST-CGG_90_-EGFP/+)* were viable. In contrast, moderate expression of the transgene *(pUAST-CGG_60_-EGFP)* produced homozygous viable flies when expressed in neurons. Therefore, we performed RT-qPCR experiments to determine the expression levels of Hsrω-n lncRNA from the heads of age- and sex-matched flies expressing *CGG_60_-EGFP* repeats and control flies (pUAST-EGFP). As shown in Figure 1 B, *Elav-*GAL4 driven *CGG_60_-EGFP* brains showed a more than three-fold increase in the level of Hsrω-n lncRNA compared to controls. In *Elav-GAL4* driven heterozygous CGG_90_/+ flies, the Hsrω-n expression resembled the two *Hsrω-n* overexpression lines: *Elav/+; Hsrω^EP93D^/+* and *Elav/+; Hsrω^EP3037^/+.* This prompted us to examine Hsrω-n RNA levels in heterozygous combinations of CGG_90_/+ and Hsrω^EP3037^/+ or *Hsrω*^EP93D^/+ flies. This combination resulted in considerable morbidity, preventing the evaluation of the expression of Hsrω-n. However, *Elav-*GAL4 driven coexpression of single copies of *CGG_60_/+* and *Hsrω^EP3037^/+* or *Hsrω^EP93D^/* + resulted in the augmentation of Hsrω-n lncRNA levels up to one and a half times compared to heterozygous EP expression alone. To investigate Hsrω-n expression at the cellular level, we used fluorescence RNA in situ hybridization (FRISH) on partially squashed salivary gland (SG) spreads, where the nuclear envelope was not disturbed. Normally, in SG chromosome spreads, Hsrω-n transcripts are detected in nucleoplasm as FRISH signals with a prominent hybridization signal at the transcription site (chromosome 3R, 93D locus) (35). We expressed CGG_60_/+ or CGG_90_/+ in the SG using the *Fkh-*GAL4 driver, which is a salivary gland-specific GAL4 driver, or a neuronal *Elav-*GAL4 driver, which also expresses in the SGs (40,41). Figure 1C shows the FRISH patterns of the Hsrω-n riboprobes in various genotypes. FRISH for cells expressing *CGG* transgene invariably showed higher Hsrω-n lncRNA expression, represented by a stronger hybridization signal and larger aggregates at the 93D site, compared to cells not expressing *CGG* repeats (Fig. 1C, columns 3 and 5 vs. columns 1). Interestingly, in SGs coexpressing single copies of *CGG_90_* and *Hsrω^dsRNA^,* the signal was weaker and more dispersed, resembling the wild-type situation (Fig. 1C, column 6). This showed that the signal aggregates were specific and not due to chromosome condensation or fixation defects. Taken together, these results suggest that Hsrω-n transcript levels are modulated by CGG-repeat length and dose.

**Figure 1:**
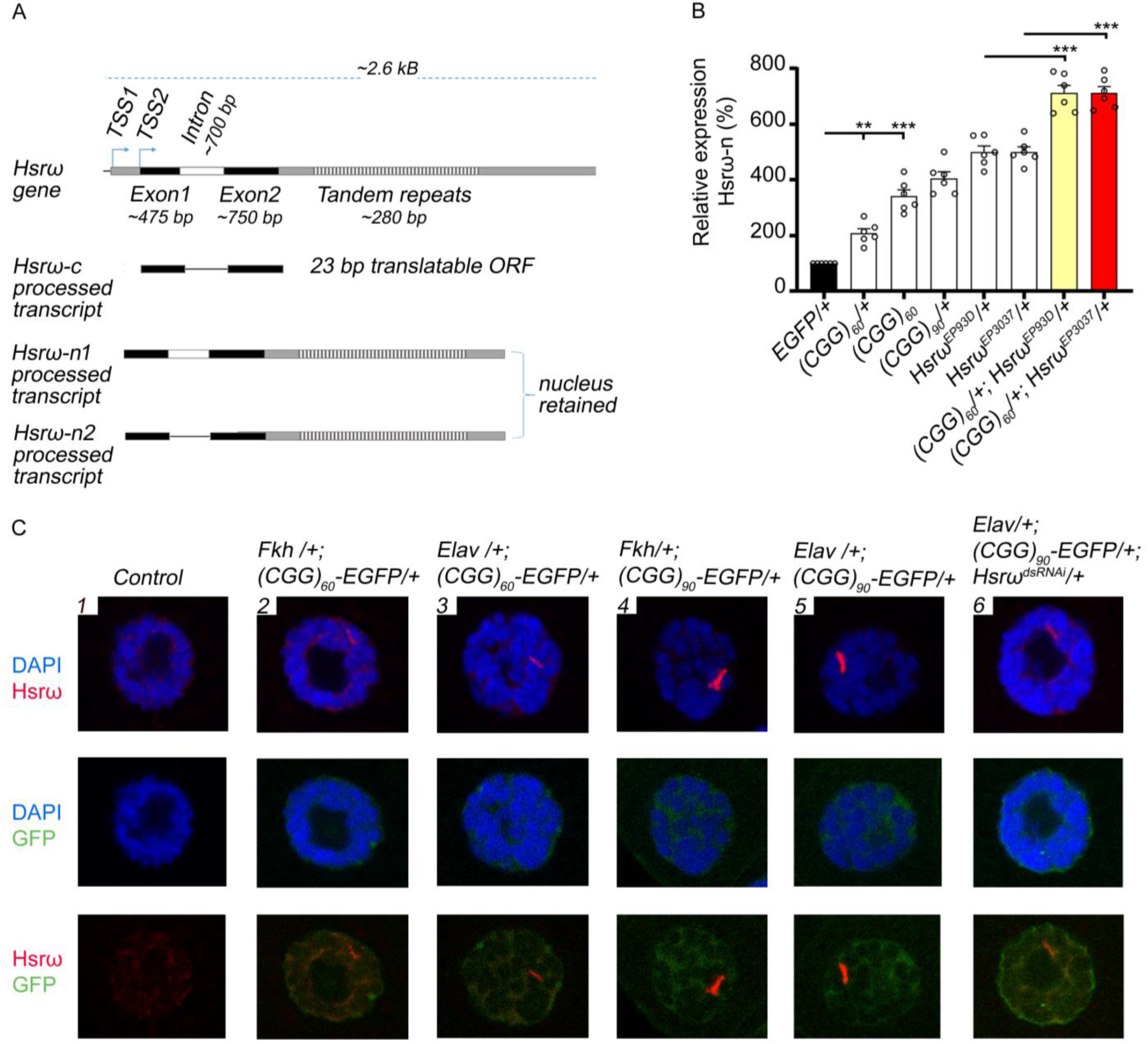
*Hsrω-n* is upregulated in CGG-repeat expressing transgenic flies. (A) Schematic representation of *the Hsrω* gene. FlyBase reports seven transcripts for the Hsrω gene; however, for simplicity, only three are shown. The spliced intron is indicated by a black line joining exons 1 and 2. (B) Relative levels of *Hsrω-n* in the adult heads of ten-day-old male flies of the indicated genotypes (bottom). *Hsrω-n* expression was increased in flies expressing CGG alone, two *Hsrω-n* overexpression lines, and combination flies compared to control flies. *** p ≤ 0.001. n ≥ 3; error bars indicate SEM. (C) Localization of *Hsrω-n* transcripts (red) in larval SG nuclei of various genotypes (top) using fluorescence RNA in situ hybridization. Chromatin is stained with DAPI (blue). Hsrω-n signal (red) in the control SG nuclei was diffused (Column 1) compared to nuclei expressing moderate CGG repeats (Column 2 and 3), or strong CGG repeats (Column 4 and 5). Repression of Hsrω-n restored the diffused appearance of the Hsrω-n signal (Column 6).

### Hsrω-n lncRNA enhances neuronal toxicity caused by expanded CGG repeats

We sought to determine the consequences of increased *Hsrω-n* RNA production in transgenic flies expressing CGG repeats. Heterozygous *Hsrω^EP3037^/+* line displayed viability comparable to that of wild-type and heterozygous *CGG_60_* flies when driven with the *Elav*-*GAL4* driver. However, viability was reduced in flies with heterozygous combinations of *CGG_60_* and *Hsrω^EP3037^* (Fig. 2A). In addition, the heterozygous combination of *Elav*-*GAL4* driven *CGG_90_* and *Hsrω^EP3037^* leads to lethality, of which 30-40% die as differentiated pupae. The remaining surviving flies showed reduced viability compared to *Elav*-*GAL4*>*CGG_60_/+; Hsrω^EP3037^/+* or *Elav*-*GAL4*>*CGG_90_/+* alone. This indicates that the overexpression of Hsrω-n lncRNA enhances CGG-mediated neuronal toxicity. Therefore, to systematically understand the consequences of increased Hsrω-n lncRNA expression in CGG repeat-mediated neurodegeneration, we used the *Drosophila* eye as a model for genetic interaction experiments, which has been extensively used in previous studies (15,20,42,43). As reported previously, in response to the *GMR-GAL4* driver, heterozygous expression of CGG_90_ produces a rough-eye morphology with disorganized ommatidia compared to the mild effect of CGG_60_ expression (15). Therefore, we used the CGG_90_ induced rough facet eye phenotype as the background for our genetic interaction experiments. We crossed the *Hsrω RNAi* line (44,45) with a recombinant line expressing CGG_90_ in the eyes *(GMR-GAL4, UAS-CGG_90_/+)* and visualized the eye phenotype of the heterozygous progeny flies *(GMR-GAL4, UAS-CGG_90_/+; Hsrω^dsRNA^/+).* As shown in Figure 2B, we observed suppression of the CGG-induced rough facet eye phenotype upon repression of *Hsrω-n* lncRNA (Fig. 2B, panel 4 vs. panel 2). In contrast, overexpression of *Hsrω-n* enhanced the phenotype in the heterozygous CGG_90_/+ genetic background (Fig. 2B, panel 6 vs. panel 2). The rough eye phenotype was extremely strong, with prominent necrotic spots along the posterior side of the eye. Notably, when driven by *GMR-GAL4,* neither Hsrω^dsRNA^ nor Hsrω^EP3037^ alone produced eye phenotypes on their own (Fig. 2B, panel 3 and 5), consistent with previous reports (46). Previously, eye degeneration by CGG transgene expression was found to be associated with the formation of inclusions or aggregates that were positive for ubiquitin, Hsp70 chaperone, and proteasome, but negative for CGG transcripts (15). Therefore, we sought to determine the effects of Hsrω-n activation on such aggregates. We performed immunohistochemistry on adult fly eye sections expressing CGG_90_ alone or co-expressing CGG_90_ and *Hsrω^dsRNA^/+.* However, we were unable to precisely determine the structure and subcellular localization of the aggregates/inclusions because of extensive degeneration/holes in the tangential sections of the eyes, which prevented us from using this phenotype in our interaction experiment. Therefore, to quantify the phenotypes of fly eyes for statistical evaluation, we manually determined phenotypic scores based on three measures: loss of pigmentation, ommatidial disruption, and necrotic spots. A higher phenotypic score represents an increased severity of the eye phenotype. As shown in Fig. 2C, modulation of the CGG_90_ induced eye phenotype by altered expression of Hsrω-n was consistent and significant across many crossings. Importantly, RT-qPCR analysis showed that a single copy of the GMR-GAL4 driver was adequate to stimulate the expression of transgenes in various genotypes (Fig. S1) Moreover, EGFP or the RAN translational product, polyG-GFP, detected by western blotting of the total protein lysate prepared from late-stage larval brains, remained unchanged in flies expressing CGG_90_ alone or in combination with *Hsrω-n* (Fig. 2D and 2E, quantification). Together, these results indicate that *Hsrω-n* modulates CGG-mediated neurodegeneration rather than altering CGG repeat transcription or RAN translation of CGG repeats.

**Figure 2:**
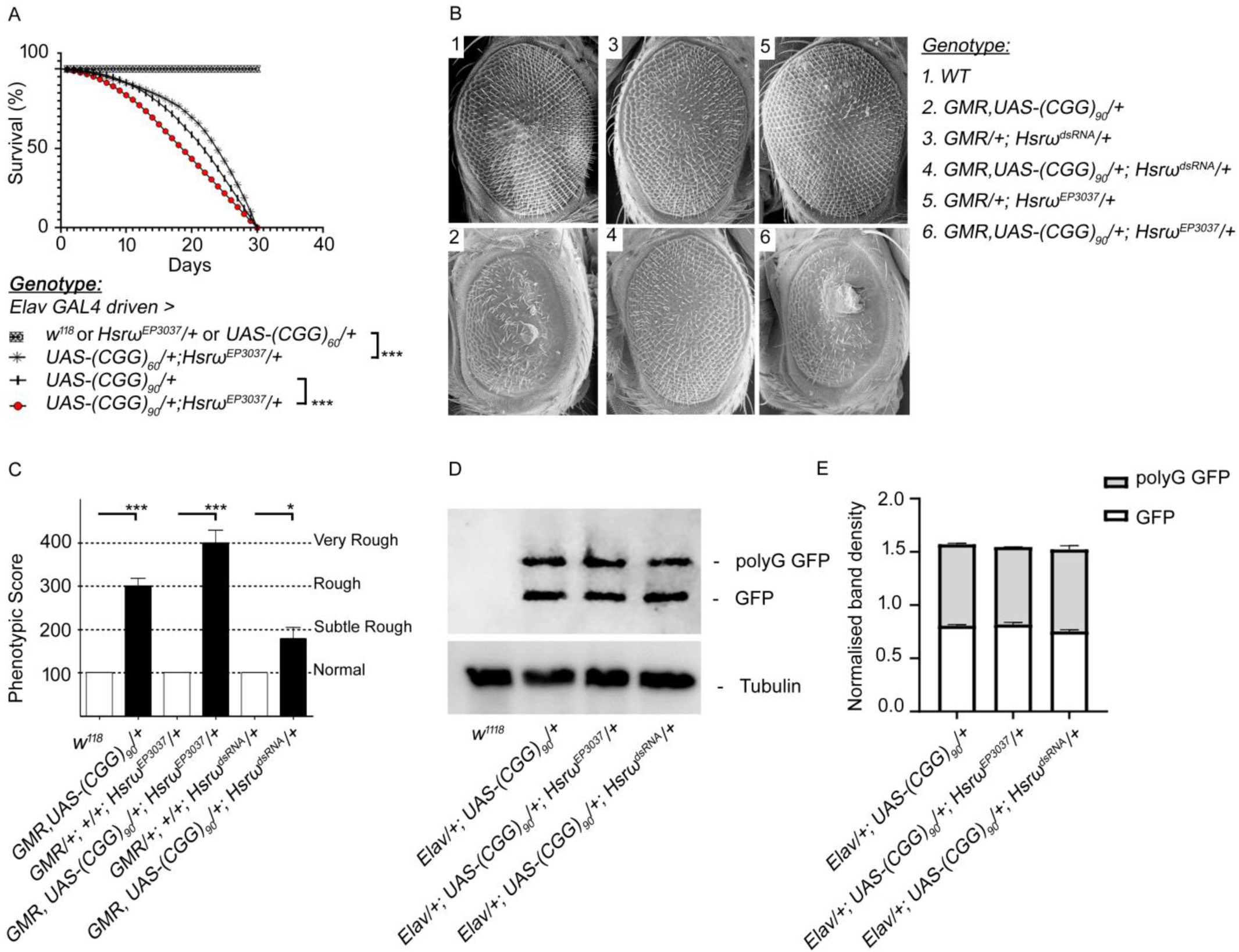
Upregulation of *Hsrω-n* expression is involved in CGG-repeat-mediated neurodegeneration. (A) Survival curves between various genotypes were analyzed by counting the surviving flies for 30 days after hatching (n=90, male adult flies) in various genotypes (bottom). Overexpression of heterozygous *Hsrω-n* significantly reduced the viability of viable *CGG_60_* flies: (*Elav/+; +/+; Hsrω^EP3037^/+) or* (*Elav/+; UAS-CGG_60_ EGFP/+),* compared with the combination flies (*Elav/+; UAS-CGG_60_-EGFP/+; Hsrω^EP3037^/+).* Similarly, compared to flies expressing stronger CGG alleles, viability was reduced in the presence of *Hsrω-n* overexpression: (*Elav/+; UAS-CGG_90_-EGFP/+)* compared to *(Elav/+; UAS-CGG_90_-EGFP/+; Hsrω^EP3037^/+).* The survival curves were significantly different (*** p ≤ 0.001). (B) Scanning electron microscopy (SEM) (1–6) of 6-8 eye images of 14-day-old flies of various genotypes as indicated. Compared to WT (1), heterozygous expression of (2) *CGG_90_* repeats results in disorganized and fused ommatidia. (3) Heterozygous knockdown or (5) overexpression of *Hsrω-n* does not elicit any eye phenotype. However, (4) heterozygous knockdown of *Hsrω-n* in the genetic background of CGG repeats rescued the neurodegenerative eye phenotype, indicating strong suppression of CGG90-mediated neurodegeneration. Similarly, (6) heterozygous overexpression of *Hsrω-n* in CGG_90_ repeat flies enhanced the rough eye phenotype. (C) Phenotypic scores (PS) based on visual assessment of the eye phenotypes of the indicated genotypes (bottom). The number of images used for these assays was n=40. The height of the histogram represents the phenotypic score (PS). A PS of 100 represented normal eye phenotypes, and a PS of 200 represented subtle rough eye phenotypes. A PS of 300 represented rough phenotypes, and a PS of 400 and above represented a very rough eye. *** p ≤ 0.001 and *p ≤ 0.05 (n ≥ 6; (n ≥ 3; error bars represent mean ± SEM). (D) Western blot depicting the RAN translation product, FMR polyG-GFP, and tubulin from late-stage larval brains expressing CGG_90_ alone or with *Hsrω^EP3037^* and *Hsrω^dsRNA^.* (E) Normalized band densities/intensities of western blot data show that *Hsrω* overexpression, *Hsrω^EP3037^* or repression, *Hsrω^dsRNA^, do not* impact *polyG* GFP and GFP protein levels in larval brains expressing CGG_90_.

### Hrb87F, the *Drosophila* homolog of hnRNP A2/B1, mediates the genetic interaction between rCGG repeats and Hsrω-n lncRNA

Hsrω-n lncRNA intricately and dynamically affects several components implicated in poly(Q) pathogenesis (46,47). Overexpression of Hsrω-n lncRNA enhances eye neurodegeneration in transgenic flies expressing expanded poly(Q) (127Q) or mutant human huntingtin protein (48). Consequently, sequestration of hnRNPs and related RNA-binding proteins by overexpression of Hsrω-n lncRNA enhances toxicity due to poly(Q) expansion (47). Interestingly, many of these proteins modulate CGG-mediated neuronal toxicity by directly or indirectly binding to the rCGG repeats. Based on this premise, we performed a genetic screen to identify the potential components through which Hsrω-n lncRNA can modulate rCGG-mediated toxicity. The screen used the *Drosophila* GAL4/UAS system to directly express CGG_90_ repeats and *Hsrω-n* simultaneously in the eye with the *GMR-GAL4* driver. This combination [*GMR, UAS-CGG_90_/+*; *Hsrω^EP3037^/+*] resulted in prominent necrotic spots along the posterior region of the eye facet (Fig. 2B, panel 6 and Fig. 3A, panel 2). The combination line was subsequently crossed with mutants of selective genes that function in the Hsrω-n lncRNA network to identify dominant modifiers (enhancers or suppressors) (Table 1). We identified Hrb87F, a homolog of human hnRNPA1, as a candidate gene that prominently suppressed the typical eye phenotype in flies expressing CGG repeats alone or in combination with Hsrω-n [*GMR, UAS-CGG_90_/+*; *Hsrω^EP3037^/+*] (Fig. 3A, column 2 vs. column 1, and Fig. 3B, quantification). Suppression was more pronounced when Hsrω-n was repressed by the Hsrω-RNAi transgene (Fig. 3A, column 3 and Fig. 3B, quantification). Conversely, heterozygous deficiency of Hrb87F in the genetic background of *GMR-GAL4, UAS-CGG_90_-EGFP*; *Hsrω^EP3037^*/+ flies led to lethality; few pharate adults had significantly damaged eyes beyond quantification, reduced eye size, almost complete loss of ommatidial arrays and bristles, and formation of black lesions on the eye surface.

**Figure 3:**
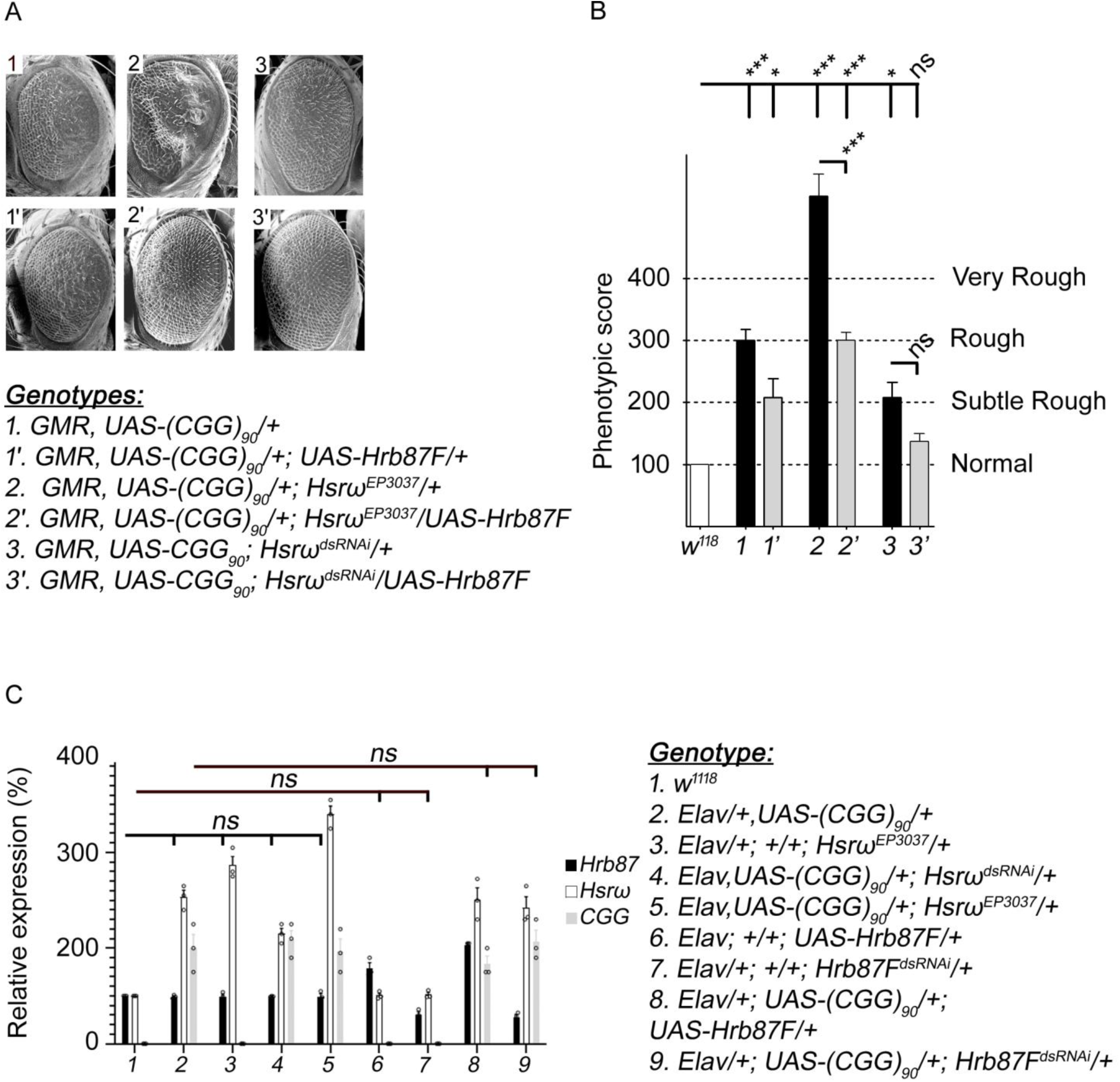
Hrb87F mediates genetic interactions between rCGG-repeats and *Hsrω-n* lncRNA. (A) SEM (6–8) eye images of 14-day-old flies with the indicated genotypes (bottom). (1-1’) Heterozygous expression of CGG_90_ repeats alone resulted in a rough eye phenotype with disorganized and fused ommatidia, which was markedly suppressed by coexpression of Hrb87F. (2-2’) Heterozygous expression of CGG_90_ repeats together with Hsrω^EP3037^ produces phenotypic readouts, typical necrotic spots, and disorganized and fused ommatidia. Coexpression of Hrb87F expression in this background suppressed the rough eye phenotype. (3-3’) Overexpression of Hrb87F in eyes co-expressing CGG_90_ and *Hsrω^dsRNA^* significantly suppressed eye degeneration. (B) Histogram indicates the quantification of interaction experiments based on phenotypic scores (PS). ***p ≤ 0.001 and ns indicates p > 0.05 (n ≥ 6; error bars represent mean ± SEM). (C) Histogram indicates the relative RNA levels of Hrb87F, *Hsrω-n*, and CGG repeats in the ten-day-old adult heads of the indicated genotypes (bottom). Overexpression of *Hsrω-n* and/or *CGG* repeats did not affect the transcription of Hrb87F. Deficiency or overexpression of Hrb87F had no effect on the transcription of CGG_90_ or *Hsrω-n* RNA. ns represents p > 0.05. (n ≥ 3; error bars represent mean ± SEM).

**Table 1:**
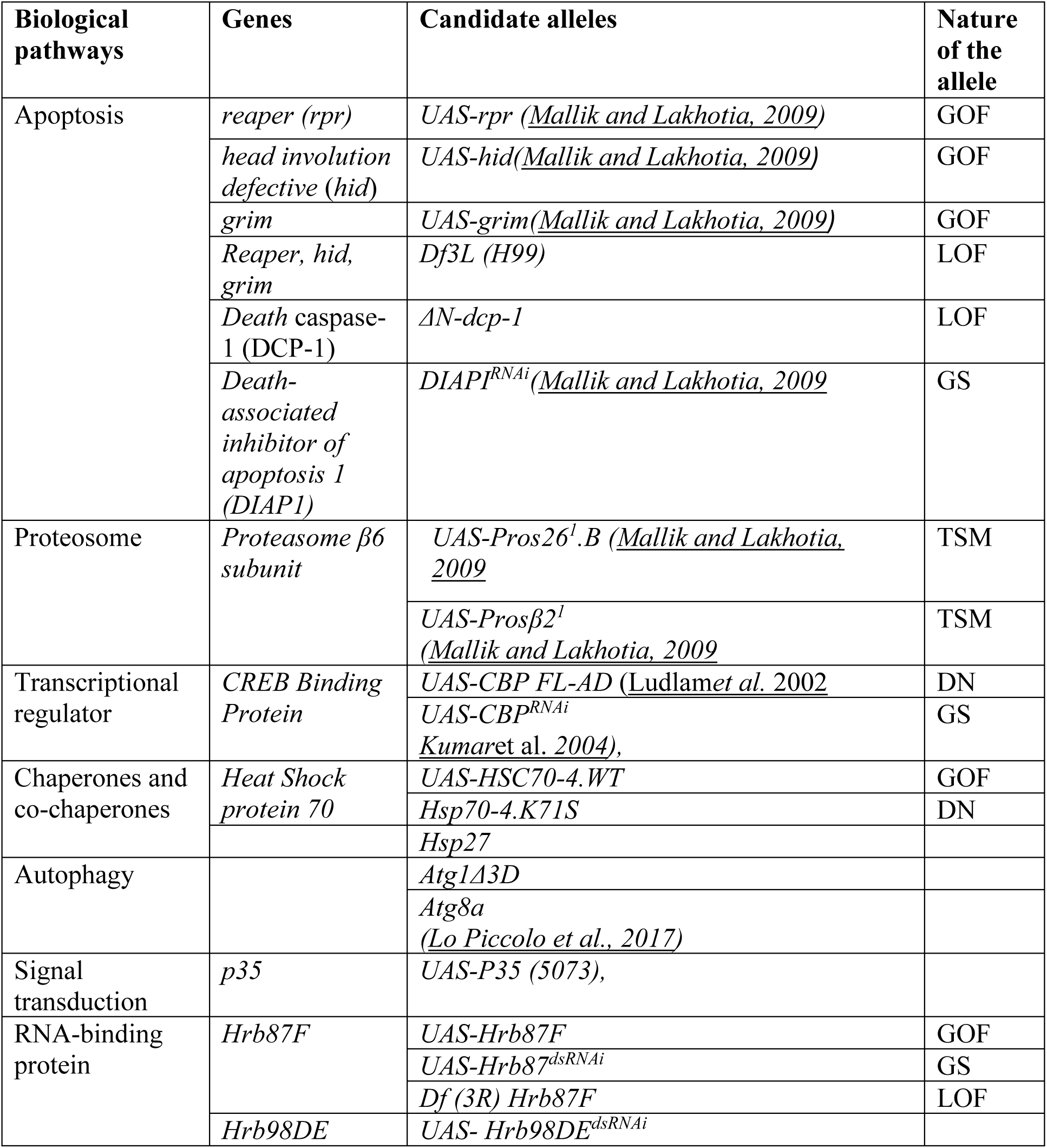
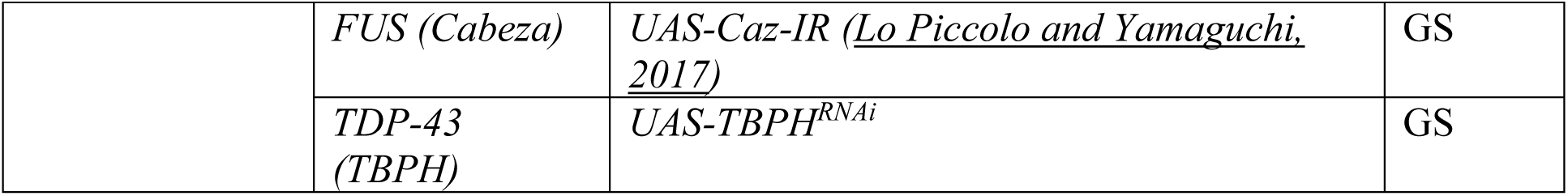
Interaction screen to identify modifiers of the eye phenotype induced by heterozygous co-expression of the (CGG)_90_ and Hsrω^EP3037^ transgenes. Column 1, Biological pathways that are directly or indirectly modulated by Hsrω-n or CGG repeat. Column 3: Alleles known to interact with Hsrω-n in modulating eye phenotypes. Column 5, effect of the candidate allele on the eye phenotype produced by heterozygous co-expression of (CGG) _90_ and Hsrω^EP3037^. False positives were eliminated by comparing heterozygous interactions separately between candidate alleles and (CGG)_90_ or Hsrω-n. The abbreviations L.O.F, GOF, TSM, and GS denote the loss of function, gain of function, temperature sensitive mutant (TSM) and gene silencing (GS), respectively.

Given the role of Hrb87F in nuclear RNA processing (49–51), we examined the transcription of Hrb87F in flies co-expressing CGG_90_ and *Hsrω-n.* Since, *Elav*-GAL4 driven co expression of CGG_90_ and *Hsrω-n* leads to adult lethality and severely compromised health conditions, we chose late-stage wandering larvae for expression analysis. As shown in Fig. 3C, heterozygous larvae co-expressing CGG_90_ or *Hsrω-n* in their larval brains did not exhibit altered Hrb87F transcription. The unaltered transcription of Hrb87F was also confirmed in flies with depleted *Hsrω-n* expression. Furthermore, neither repression nor overexpression of Hrb87F affected CGG_90_ or *Hsrω-n* transcription. Taken together, these observations suggest that Hrb87F modulates the genetic interactions between rCGG repeats and Hsrω-n RNA at the post-transcriptional level.

### Expanded CGG repeats increased the biochemical association between *Hsrω-n* and Hrb87F

Previous studies have shown that Hrb87F physically and genetically interacts with Hsrω-n lncRNA (47), as well as with rCGG repeats (20). Given our results showing that Hrb87F mediates the genetic interaction between CGG repeats and *Hsrω-n* lncRNA, we wondered whether CGG-repeat expression affects the association of *Hsrω-n* lncRNA with Hrb87F (21,52). We conducted a cross-linking RNA-immunoprecipitation (CLIP-RIP) biochemical assay using anti-P11, an Hrb87F-specific antibody, on larval brains expressing either (1) EGFP only (control) and (2) CGG_90_-EGFP. As a control for non-specific associations, the same experiments were performed with IgG antibodies in parallel. Following Clip-RIP, immunoprecipitated material was quantified for CGG and Hsrω-n transcripts, as well as two other abundant nuclear non-coding RNAs, Rox1 and U4 that served as specific controls. Normalization of the amount of each RNA detected in the IP samples with the corresponding RNA amount in the input revealed a significant presence of Hsrω-n RNA with Hrb87F in the presence of CGG repeats (Fig. S2). As a positive control, the levels of CGG RNA, a well-validated target of Hrb87F, were also increased. Fold change in the enrichment values of the normalized values (IP^P11^/Input over IP^IgG^/Input) is significantly higher in presence of CGG repeats for Hsrω-n and CGG transcripts, while Rox1and and U4 transcripts show minimal differences between control and in presence of CGG repeats (Fig. 4A). The fold difference values also revealed that compared to Rox1 (+0.059-fold) and U4 (-0.034-fold), Hsrω-n associates more with Hrb87F in the presence of CGG repeats (+2.864-fold). Overall, these findings support the possibility that Hrb87F binding with Hsrω-n is enhanced in the presence of CGG repeats Considering the enrichment of Hsrω-n lncRNAs with Hrb87F in the presence of CGG repeats, we speculated that Hrb87F also showed an altered distribution in the presence of CGG repeats. To answer this question, we conducted immunofluorescence on polytene chromosomes derived from squashed SG nuclei. Normally, Hrb87F decorates transcriptionally active interband region of polytene chromosomes (43) (Fig. 4B, column 1). However, in the presence of CGG repeat expression, most of them disappeared from their normal chromosome sites, except for the 93D band at chromosome 3, which is the site of *Hsrω* gene (53). Because Hsrω-n scaffolds the formation of the ω-speckle together with hnRNPs at the 93D site during stressful conditions, we sought to determine Hrb87F binding to nascent Hsrω-n RNA at the 93D site. However, due to technical difficulties with our immuno-FRISH experiments, we could not show direct binding of Hrb87F to nascent Hsrω-n lncRNAs at the 93D site. Nonetheless, with the expression of the stronger CGG allele, the Hrb87F at 93D site appeared intensely aggregated compared to moderate expression of the CGG allele (Fig. 4B, Column 3 vs. Column 2), suggesting redistribution of the Hrb87F protein in the presence of CGG repeats.

**Figure 4:**
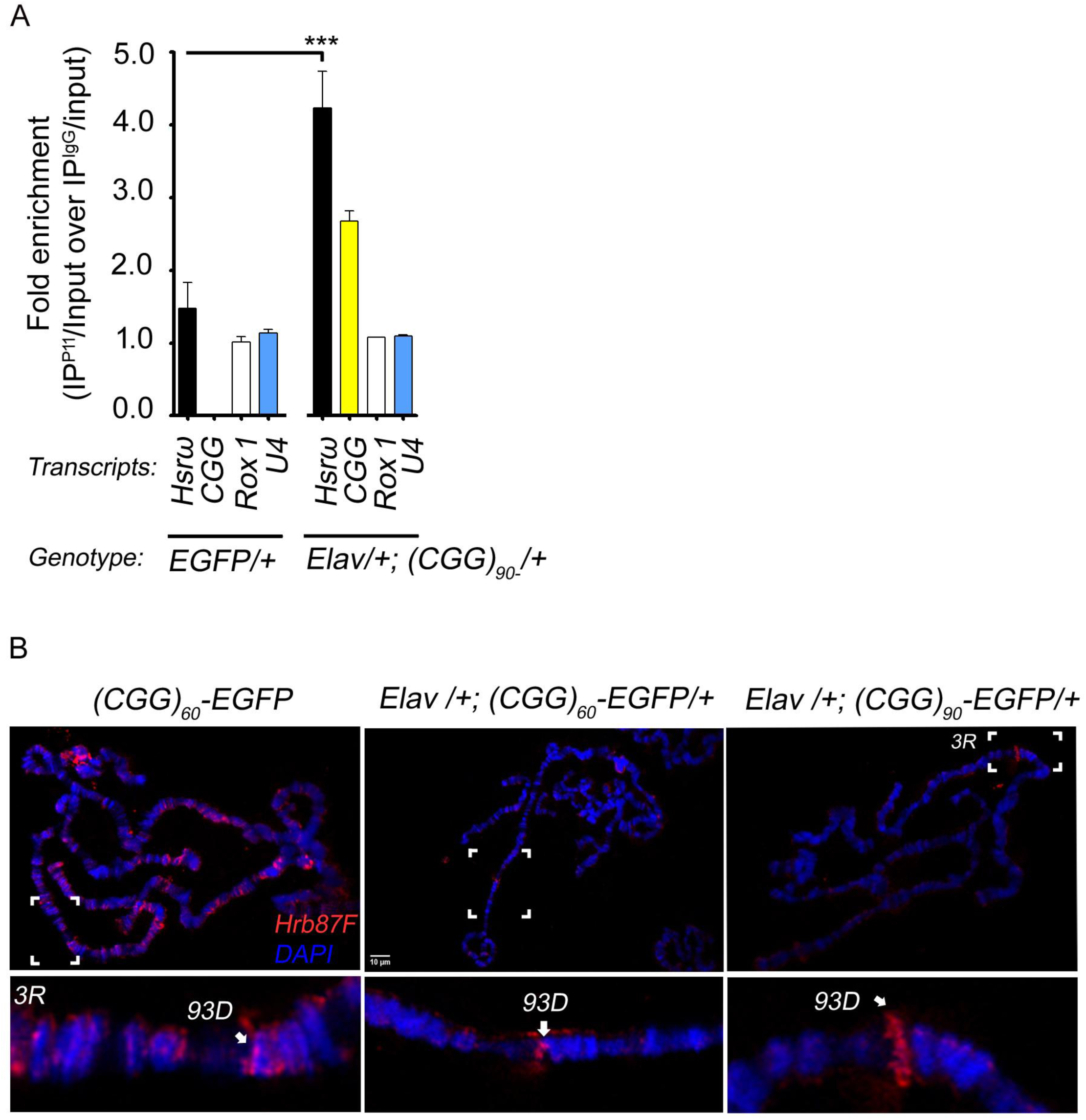
Hrb87F biochemical association with *Hsrω-n* lncRNAs is enriched in the presence of rCGG repeats. Clip-RIP and subsequent RT-qPCR on larval brains from the indicated genotypes (bottom). Following Clip-RIP, the Input (10%), P11, and IgG-immunoprecipitated complexes from the indicated genotypes (bottom) were used for the RT-qPCR analysis. For each gene under study, the amount of RNA detected in the IP samples was normalized to the amount of the corresponding gene in the input (IP^P11^/input and IP^IgG^/input) (A) The Histogram represents the fold enrichment in the expression of target genes present in the P11 immunoprecipitated RNA divided by the corresponding expression present in the input lysates normalized to the same ratio in the IgG control Clip-RIP. Significantly more Hsrω-n was bound to P11 (Hrb87F) in the presence of the CGG repeat. *** p ≤ 0.001 (n ≥ 3; error bars represent mean ± SEM). Differences in the levels of Rox1 and U4 were not statistically significant. (B) Confocal projections of Hrb87F (red) and DAPI-stained (blue) polytene chromosomes from larval SG of the indicated genotypes (top). Magnified views of chromosome 3R are shown in parallel (bottom panel). Arrows indicate the accumulation of Hrb87F (red) in the 93D puff on the 3R chromosomal arm. Chromosome 3, 93D site was determined in reference to the standard SG polytene chromosome pattern (66). In the presence of CGG repeats, Hrb87F showed increased accumulation at the 93D locus compared to control polytene chromosomes not expressing CGG repeats.

### Hsrω-n lncRNA downregulation restored nuclear Hrb87F and cytoplasmic Hsp70 protein distribution in CGG repeat expressing cells

CGG repeats are known to sequester a large fraction of RBPs such as hnRNP A2/B1 and Pur α, which can disrupt RNA processing (20,21). Previously, we showed that CGG repeats cause upregulation of Hsp70 mRNAs that are paradoxically retained within the nucleus instead of being turned over to the Hsp70 protein (32). Since ω-speckles play essential roles in the storage/sequestration of the hnRNP family and other proteins during stress (35), we wondered whether the dominant suppression of CGG-induced degeneration by Hsrω-n RNAi affected hnRNP or Hsp70 protein distribution. We examined the distribution of Hrb87F and Hsp70 in partially squashed SGs from late third-instar larvae expressing CGG repeats alone or in combination with Hsrω-RNAi. In the presence of expanded CGG repeats, Hrb87F exhibited an aggregated pattern at the 93D locus (Fig. 5A, column 2). In contrast, the Hsp70 protein exhibited a diffuse distribution around the nucleus with no statistically significant increase in protein levels (Fig. 5B, quantification), despite a three-fold increase in Hsp70 mRNA levels (Fig. 5C, column 2, quantification). This is in line with our previous studies that showed nuclear accumulation of Hsp70 transcripts due to nuclear export defects (32). Notably, co-expression of Hsrω-RNAi and CGG transgenes resulted in the diffusion and relocalization of aggregate-associated Hrb87F to the nucleoplasm (Fig. 5A, column 3 and 5B, quantification). Interestingly, this was also associated with a significant presence of Hsp70 in the cytoplasm in a diffuse pattern around the nucleus (Fig. 5A, column 3 and 5B, quantification). It is worth noting that Hsrω-RNAi did not alter the transcriptional status of Hsp70 or Hrb87F (Fig. 5C and 3C). These data suggest that the loss of Hsrω-n lncRNA releases RBPs associated with ω-speckle, which enhances the titer of functionally available RBPs in the cell, consequently facilitating proper RNA processing or nuclear transport and rescuing the disease phenotype. Previously, ectopic expression of Hsp70 has been shown to suppress the cellular toxicity associated with CGG repeats (15). Based on the data provided above, alterations in Hsp70 may be a consequence rather than a cause of reduced CGG neurotoxicity.

**Figure 5:**
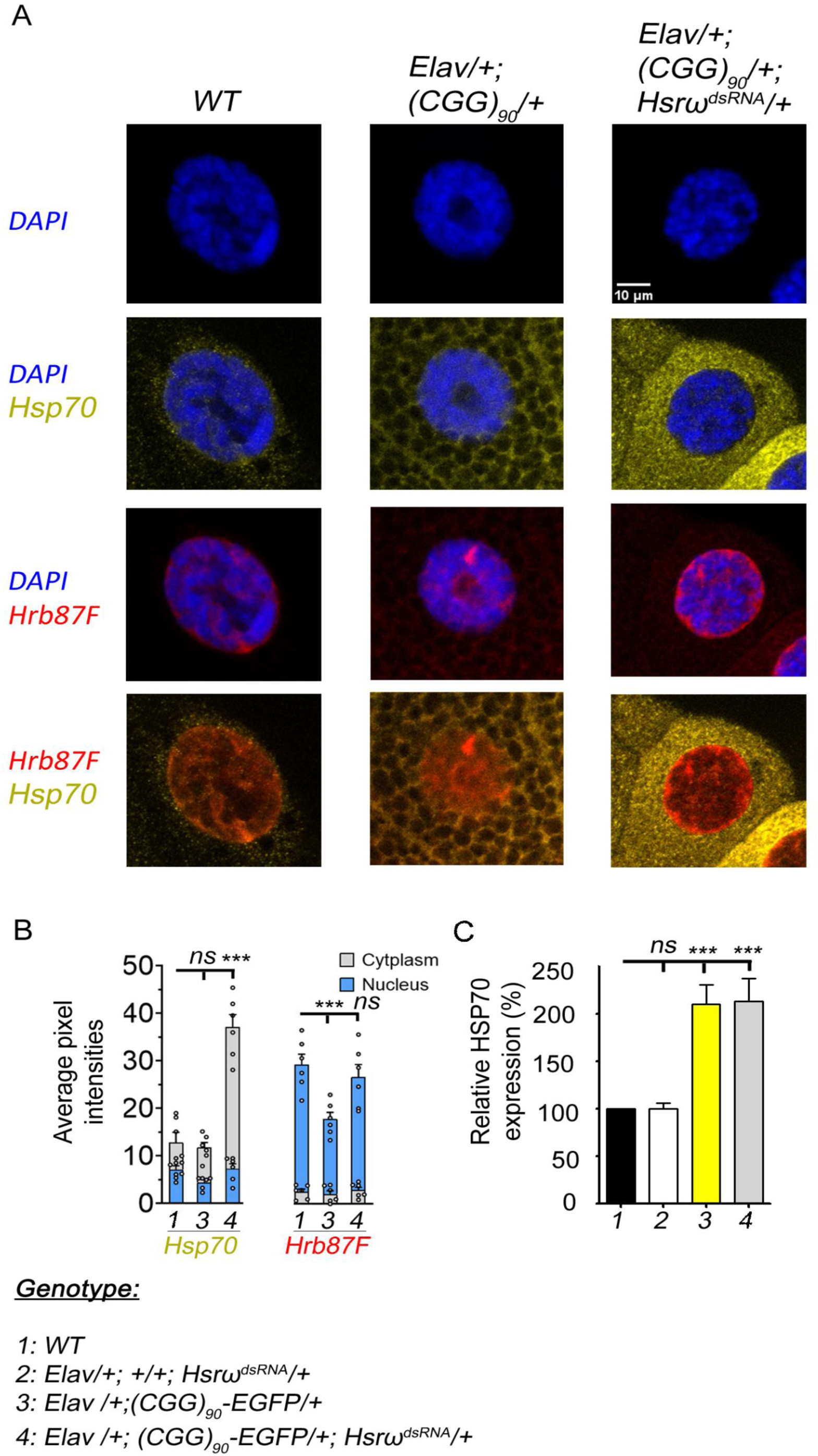
Hrb87F accumulates at the 93D chromosomal locus in the presence of CGG-repeats. (A) Confocal projection images showing immunolocalization of Hsp70 (yellow) and Hrb87F (red) in the SG nucleus of the indicated genotypes (top). DNA was stained with DAPI (blue). SGs expressing CGG repeats showed diffused appearance of Hsp70 (yellow) around nucleus resembling wildtype SGs. However, Hrb87F appeared aggregated in the nucleus compared to the control. Downregulation of Hsrω-n transcripts in SGs expressing CGG repeats caused enhanced diffusion of Hsp70 (yellow) around nucleus. Downregulation of Hsrω-n transcripts in SGs expressing CGG repeats also caused aggregated Hrb87F to diffuse and relocalize to the nucleoplasm. (B) Histograms illustrate the percentage of average pixel intensities associated with Hrb87F or Hsp70 within the nuclei and cytoplasm of the specified genotypes. Statistical significance is denoted as follows: ***p for Cytoplasm (Hsp70) ≤ 0.001; ^ns^p for Cytoplasm (Hsp70) > 0.05; ***p for Nucleus (Hrb87F) ≤ 0.001; ^ns^p for Nucleus (Hrb87F) > 0.05. In all cases (n ≥ 60 nuclei for each genotype corresponding to 6 larvae; error bars represent mean ± SEM). (C) Quantitative analysis of *Hsp70* mRNA levels by real-time PCR from the larval brains of the indicated genotypes (bottom). Hsp70 RNA was significantly expressed in the presence of CGG repeats. *** p ≤ 0.001. n ≥ 3; error bars indicate SEM.

### Expanded CGG repeats activate *SatIII* and *Neat 1* expression in mammalian cells

In mammalian cells, nuclear stress bodies (nSBs) and paraspeckles are functional analogs of ω-speckles and share the requirements for their formation. They are built on specific arcRNA molecules, which include highly repetitive satellite III (*SatIII*) for nSbs and nuclear paraspeckle assembly transcript 1 (*Neat 1*) for paraspeckle formation, along with protein components that include several RNA-binding proteins and SWI/SNF chromatin-remodeling complexes (54). Therefore, we investigated whether *SatIII* and *Neat I* were misregulated in mammalian cells in the presence of CGG repeats. Previously, pcDNA3.1 mammalian expression vectors containing CGG repeats of various lengths or FMR1 5’-UTR have been used in several studies (13,22,55,56). Using the same constructs, we performed transient transfection in human embryonic kidney 293T (HEK293T) cells. However, such transfections did not form RNA aggregates in cell culture, which is in line with several previous studies (13, 29). Therefore, we performed an MTT assay to determine CGG-induced toxicity in cell culture. In cell culture, CGG repeats induced toxicity three days post-transfection (Fig. 6A). Interestingly, RT-qPCR showed an almost three-fold increase in *Sat III* and a four-fold increase in *Neat I* expression in total RNA isolated from cells expressing CGG repeats compared to that in non-transfected cells (Fig. 6B). No significant change was detected in the unrelated GAPDH control (Fig. 6B). Finally, we analyzed the transcriptomics of a FXTAS mouse model that expresses PM-length CGG RNA as well as the FMRpolyG protein (27). In these mice, the age of onset was three months, and the mice were considered to have full phenotypes by six months. A comparison of the cerebellum of six-month-old and three-month-old mice showed a clear correlation between the upregulation of *Neat 1* and disease progression in both males and females (Fig. 6C). Unfortunately, we could not analyze *Sat III* expression using the transcriptomic data of these mice due to the large number of loci with *Sat III*.

**Figure 6:**
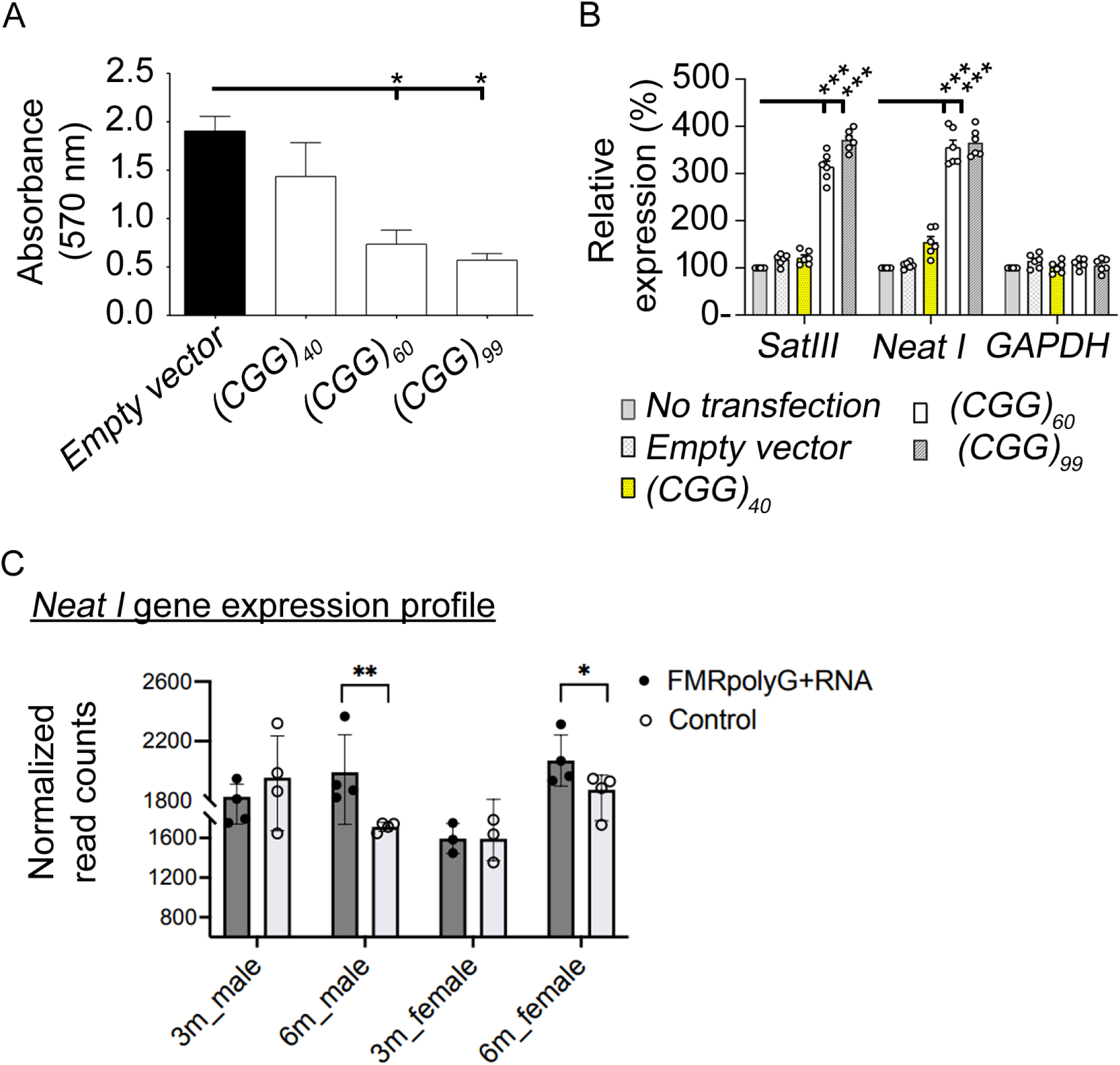
*Sat III* and *Neat I* are upregulated in cells transiently expressing CGG repeats of various lengths. Cell viability following transient expression of CGG-repeats in HEK293T cells. (A) Histogram showing cell viability in HEK293T cells transiently expressing the indicated constructs (bottom). Compared to cells transfected with the empty vector, cells transfected with CGG_40_, CGG_60,_ and CGG_99_ repeats showed significant toxicity. *p ≤ 0.05. n ≥ 3; error bars indicate SEM. (B) RT-qPCR analysis of *Sat III, Neat 1*, and GAPDH levels in untransfected and transfected HEK293T cells. Shown are relative RNA levels in cells transiently expressing EGFP, CGG_40_, CGG_60_, or CGG_99_ repeats after 3-day of transfection. The relative RNA levels were normalized to *β-actin* mRNAs levels. *** p ≤ 0.001 (n>3; error bars indicate mean ± SEM). (C) Gene expression levels of *Neat1* in transcriptomic data from cerebellum of FXTAS transgenic mice and controls. *p ≤ 0.05, **p < 0.005 from differential expression analysis of transcriptomic data.

## Expanded CGG repeats increased the biochemical association between *Hsrω-n* and Imitation SWI (ISWI), SWI/SNF chromatin-remodeling factor

Given that the SWI/SNF chromatin remodeling complex is an important structural and common subunit in ω-speckles, nSBs, and paraspeckles (34), we hypothesized that in the presence of CGG repeats, overexpression of *SatIII* or *Neat I* lncRNAs could lead to increased recruitment of SWI/SNF to the respective nuclear bodies, which subsequently affects the transcription and/or RNA processing of the respective arcRNAs. In *Drosophila*, Imitation SWI (ISWI), a highly conserved member of SWI2/SNF2, forms the catalytic center of a wide variety of chromatin-remodeling complexes (57,58). In addition, it was identified as a factor that physically and genetically interacts with *Hsrω-n* lncRNAs to form ω-speckles (43). Therefore, we sought to determine whether ISWI was redistributed in ω-speckles in the chronic presence of expanded CGG repeats. We performed two series of transfections in *Drosophila* S2 cells. In the initial series, we independently evaluated the toxicity induced by transfection with either His-tagged ISWI or CGG_99_ repeats. In a subsequent transfection, His-tagged ISWI and CGG_99_ repeats were transiently co-expressed. Notably, the expression of CGG_99_ repeats after three days of transfection was associated with cellular toxicity in culture. Importantly, the co-expression with His-tagged ISWI after three days did not alter the toxicity associated with CGG repeat expression (Fig. 7A). The CLIP-RIP assay was therefore performed on the indicated transfections after three days, using anti-His and generic IgG antibodies followed by RT-qPCR against CGG or *Hsrω-n*, and the specificity controls Rox1 and U4 (Fig.7B). Normalization (IP/input) of the amount of each RNA under study in the IP samples with the corresponding RNA in the input samples showed significant enrichment of Hsrω-n RNA in cells co-transfected with His-tagged ISWI and CGG_99_ compared to His-tagged ISWI expression. Accordingly, fold change in the enrichment values of the normalized values (IP^His-ISWI^/Input over IP^IgG^/Input) showed a 1.921-fold increase in Hsrω-n RNA immunoprecipitated with His-ISWI in the presence of CGG repeats (Fig 7B). However, no significant enrichment was detected for Rox1 or U4, suggesting that ISWI protein associates with Hsrω-n lncRNA. Notably, as shown in Fig. 7C and 7D, quantification, ISWI protein abundance was unchanged across input samples with or without CGG repeats. In addition, mRNA levels of hSNF2h/ISWI in HEK293T cells with CGG repeats remained unchanged (Fig. 7E). Therefore, differences in Hsrω-n lncRNA recovery in CLIP-RIP assays with CGG repeats are unlikely to result from altered levels of ISWI and instead suggest for possible differences in Hsrω-n lncRNA –His-ISWI associations, the functional consequences of which remain to be determined.

**Figure 7:**
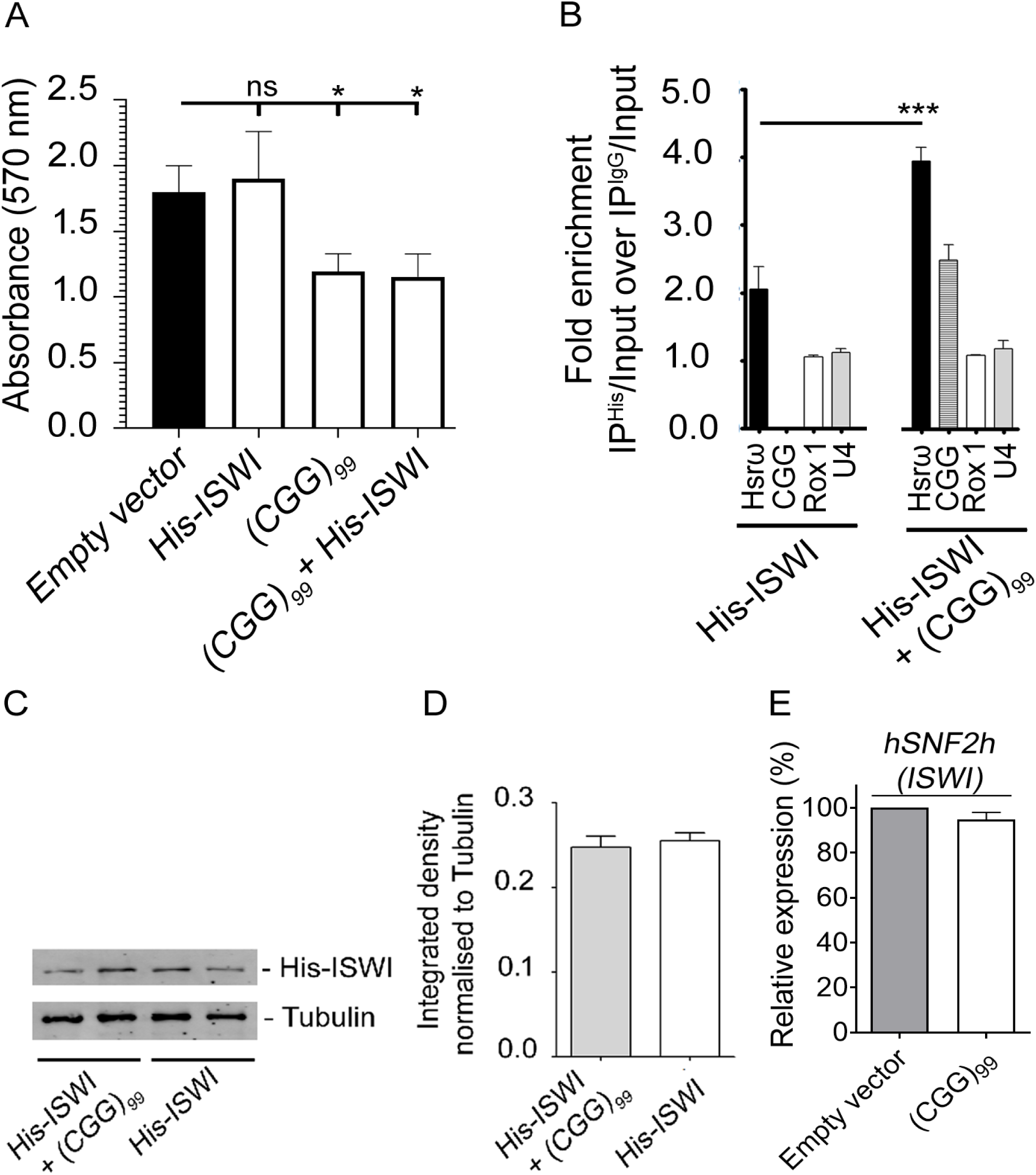
ISWI biochemical association with *Hsrω-n* lncRNAs is enriched in the presence of rCGG-repeats. (A) Cell viability following transient expression of CGG-repeats in S2 cells. Histogram showing the viability of S2 cells transiently expressing the indicated constructs (bottom). In comparison with controls, cells expressing *CGG_99_* repeats or cells co-expressing *CGG_99_* repeats and His-ISWI, showed significant toxicity. * p ≤ 0.05 (n ≥ 3; error bars indicate SEM). (B) Clip-RIP experiments were performed using an anti-histidine antibody from crosslinked S2 cells transfected prior with indicated constructs (bottom). The input (10%), anti-histidine, and IgG-immunoprecipitated complexes were used for RT-qPCR. (B) Histogram represents the fold change of indicated RNAs present in the IP samples over the corresponding expression present in the input samples normalized to the same ratio in the IgG control Clip-RIP (fold enrichment values). Significantly more *Hsrω-n* was bound to His ISWI in the presence of the CGG repeat. *** p ≤ 0.001 (n ≥ 3; error bars indicate SEM). Differences in the levels of Rox1 and U4 controls, were not statistically significant. (C) Western blotting depicting His-ISWI and tubulin from transient transfections of the indicated constructs. (D) Normalized band densities/intensities show that His-ISWI protein level remains same in presence or absence of CGG repeats. (E) RT-qPCR analysis showing the relative expression of hSNF2h (ISWI) in HEK293T cells transfected with CGG_99_ repeats compared to non-transfected cells.

### ISWI deficiency suppresses CGG repeat-induced eye phenotype

Due to the lack of specific antibodies, we were not able to determine the subcellular distribution of ISWI and therefore focused on understanding its functional role. We have shown that expression of expanded CGG repeats leads to the accumulation of hnRNPs, most notably Hrb87F, at the 93D locus (which is the site of Hsrω-n synthesis and ω-speckle formation), where they form prominent nuclear aggregates (Fig. 5A). ISWI is required for the proper aggregation of hnRNPs at the Hsrω gene locus (72). Although not all speckles visibly overlap with ISWI due to their high mobility, transient contact appears essential for speckle assembly. We asked whether ISWI reduction might influence CGG-induced toxicity. To test this, we performed genetic interaction experiments. Partial reduction of ISWI significantly suppressed the rough eye phenotype caused by CGG repeat expression, resulting in a marked rescue of eye morphology (Fig. 8A, column 3 vs. column 2, and Fig. 8A, quantification). Notably, mRNA levels of ISWI in fly brains with CGG repeats or over expressing Hsrω-n remained unchanged (Fig. 8B), suggesting a post-transcriptional role for ISWI. ISWI is well known for its functions in chromatin structure and histone H1 incorporation, with broad effects on gene expression. Accordingly, we asked whether ISWI expression modulates the transcription of Hsp70 and Hsrω-n RNA that were seen up regulated in the presence of CGG repeats. However, transcription of Hsrω-n lncRNA, or Hsp70 mRNA along with the control Hrb87F was unchanged across larval brains expressing CGG repeats with and without partial knockdown of ISWI. Thus, Hsrω-n lncRNA appears to functionally interact with ISWI and potentially influence its activity without a reciprocal change in transcript abundance when ISWI amount is altered. Taken together, these genetic and functional observations support a mechanistic model in which expanded CGG repeats increase the expression of stress-related arcRNAs, impacting the distribution of hnRNPs and/or SWI/SNF complexes to influence transcription and/or RNA processing (Fig. 9).

**Figure 8:**
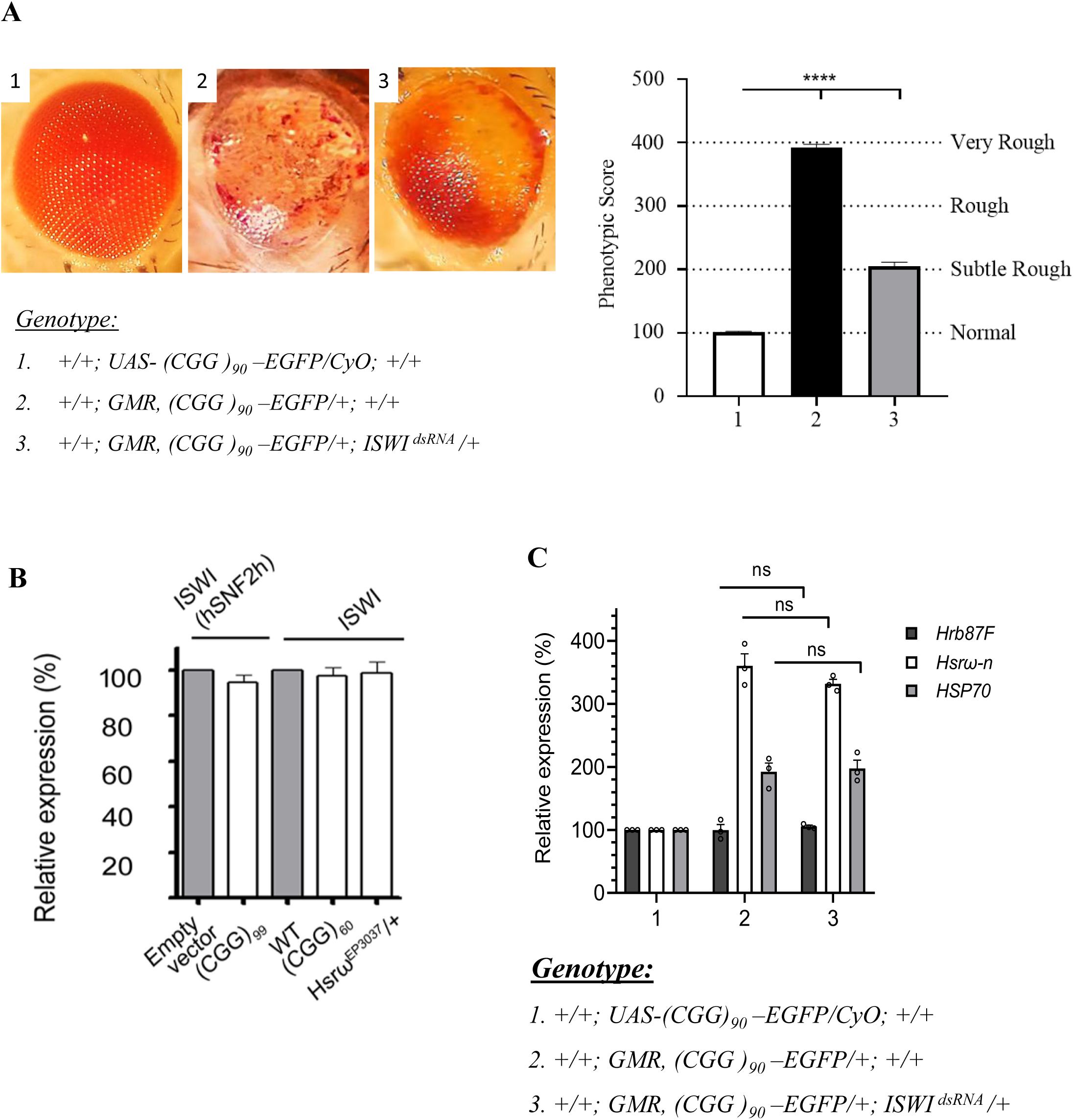
ISWI deficiency suppresses CGG repeat-induced eye phenotype. (A) Stereoscopic eye images of 14-day old flies with the indicated genotypes (bottom). Compared to the (1) Wild type flies, (2) heterozygous expression of CGG repeats results in a rough eye phenotype with disorganized and fused ommatidia. (3) Heterozygous repression of ISWI in the genetic background of CGG_90_ repeats rescued the rough eye phenotype, indicating suppression of CGG repeat-mediated neurodegeneration. (B) Quantification of eye phenotypes of 14-day old flies with the indicated genotypes (bottom). The height of the histogram represents the phenotypic score (PS). A PS of 100 represented normal eye phenotypes, and a PS of 200 represented subtle rough eye phenotypes. A PS of 300 represented rough phenotypes, and a PS of 400 and above represented a very rough eye. ****p ≤ 0.0001, n ≥ 40; error bars represent mean ± SEM). (C) RT-qPCR analysis showing the relative expression of ISWI in transgenic flies expressing expanded CGG_60_ repeats or Hsrω-n in the brain compared to WT flies. (D) RT-qPCR analysis showing the levels of indicated transcripts in the larval brains expressing (2) CGG_90_ repeats alone, and in the larval brains (3) co-expressing CGG_90_ repeats and ISWI^dsRNA^ ^ns^p > 0.05, n=3, error bars indicate the mean SEM.

**Figure 9:**
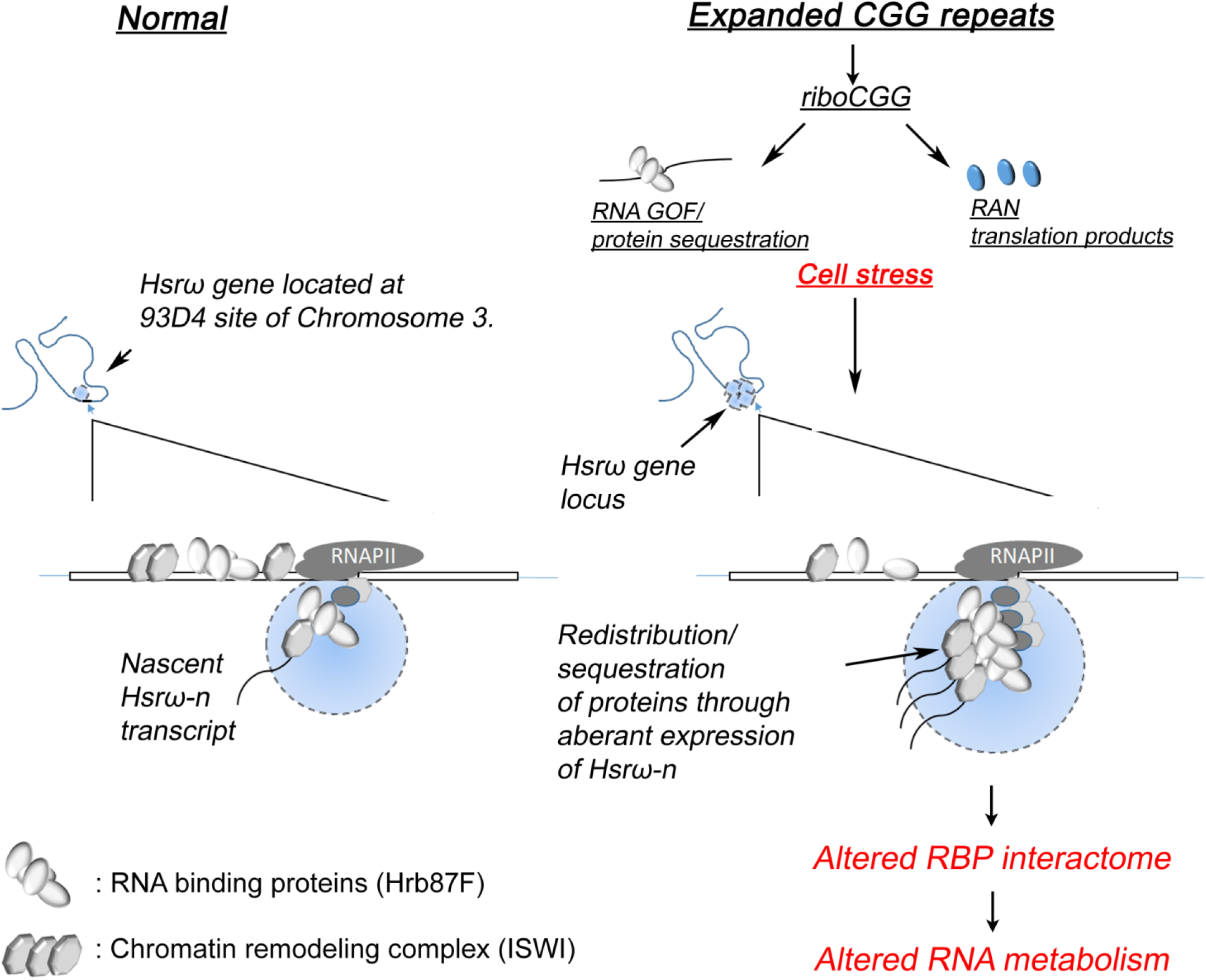
Proposed model of Hsrω-n lncRNA modulating CGG repeat-mediated neurodegeneration. Compared to normal cellular processes (left), the expression of CGG repeats could provide chronic stress, causing stress-related lncRNAs to be overexpressed. The *Hsrω-n* lncRNA, which scaffolds nuclear bodies known as ω-speckles, is shown here. Overexpression of *Hsrω-n* lncRNA can redistribute or sequester Hrb87F RNA-binding proteins and potentially ISWI chromatin-remodeling factor from their normal cellular functions, resulting in altered genomic stability and cellular homeostasis associated with FXTAS.

## Discussion

FXTAS is a late-onset neurodegenerative disorder due to 55-200 CGG repeats, referred to as PM alleles, in the 5′ untranslated region (5′ UTR) of the FMR1 gene (6). The pathogenic mechanisms include elevated levels of rCGG repeats transcribed from the expanded PM allele (59). Expanded rCGG repeats are thought to elicit RNA gain-of-function toxicity by sequestering essential RNA-binding proteins (RBPs) from their normal functions or undergoing repeat-associated non-AUG (RAN) translation to produce toxic homopolymeric peptides such as FMRpolyG (22–27,60). The effects of these mechanisms include dysregulation of multiple and diverse cellular pathways that consequently lead to chronic cellular stress and cell death. Normally, cells respond to stress by activating mechanisms that maintain homeostasis to support cellular functions. Regulatory RNAs are part of genetically encoded response networks to a variety of stress responses (61). In this study, we report that aberrant regulation of stress-inducible long non-coding RNAs can contribute to CGG-mediated toxicity in *Drosophila* and mammalian systems. Our biochemical and genetic studies demonstrated that expanded CGG repeats in flies lead to the activation of *Hsrω-n* expression.

*Hsrω-n* belongs to a class of lncRNAs referred to as architectural RNAs (arcRNAs) that function as scaffolds for nuclear bodies, termed omega speckle compartments (ω-speckles), on which protein complexes assemble to regulate gene expression (34). Under normal conditions, ω-speckles are observed in varying numbers in interchromatin regions to regulate the trafficking and availability of hnRNPs and other related RNA-binding proteins in the cell nucleus (53). However, under stressful conditions, the levels of Hsrω-n lncRNA increase at its gene locus, which possibly causes a greater sequestration of hnRNPs to protect against illegitimate RNA-processing activities. As the cells recover from stress, the normal levels of Hsrω-n lncRNA are restored, and consequently, hnRNPs and other RNA-binding proteins are released from the *Hsrω* gene locus to carry out their normal biological functions.

Many ncRNAs involved in systematic stress responses have been described in mammals (61). For example, the expression of stress-induced satellite III repeat RNAs of varying lengths, *Sat III* lncRNA, is elevated to form scaffolds for specific transcription factors and RNA-binding proteins termed nuclear stress bodies (nSBs) (62). Several proteins, including hnRNPs, were shown to be associated with *Sat III* lncRNA in nSBs after heat shock, similar to the restriction of proteins to ω-speckles in heat-shocked cells. *Sat III* also binds to TDP-43 and promotes its elongation by binding to the ELL2 domain during transcription, which can affect TDP-43 neurotoxicity (63,64). Similarly, paraspeckles are nuclear condensates built on the long non-coding RNA *Neat 1*. *Neat 1* RNA molecules may sequester hundreds of proteins and are transcriptionally upregulated in response to cellular stresses and disease states such as amyotrophic lateral sclerosis, frontotemporal lobar degeneration, and prostate and breast cancers (34,62). Although *Hsrω-n* does not share sequence similarity with *Sat III* or *Neat 1* lncRNAs, based on their functional features and regulatory mechanisms, it would be interesting to investigate whether these lncRNAs may become dysregulated in FXTAS patients. Consistent with this, we showed that *SatIII* and *Neat I* in mammalian cells are activated in the presence of expanded CGG repeats and potentially contribute to an active role in CGG-repeat-mediated neurodegeneration.

Given the role of these lncRNAs in combating stress, it is surprising that the Hsrω-n lncRNA dominantly enhanced expanded CGG repeat toxicity. Because of the chronic stress caused by CGG repeats, Hsrω-n lncRNA levels remained consistently high, which affects homeostasis of its binding partner such as hnRNP and several other proteins. Sequestration of hnRNPs by overexpression of *Hsrω-n* also seems to cause polyglutamine-induced neurodegeneration (42,47). Furthermore, *Hsrω-n* was linked to the dysregulation of other hnRNPs, including dFUS and TDP-43. These proteins are associated with the pathogenesis of amyotrophic lateral sclerosis (ALS) and frontotemporal lobar degeneration with ubiquitinated inclusions (FTLD-TDP) (65).

As *Hsrω-n* and other arcRNAs sequester specific RNA-binding proteins, thereby affecting their ability to regulate transcription and/or RNA processing, synergistic sequestration of Hrb87F by rCGG repeats and *Hsrω-n* may contribute to genomic instability and neurotoxicity. Pertinent to this, in flies, the activation of several retrotransposons in the presence of expanded CGG repeats has been attributed to the altered binding of hnRNP A2/B1 with the genomic regions containing retrotransposons (51). Similarly, SWI/SNF, which forms the catalytic center of a wide variety of chromatin-remodeling complexes, has a well-characterized role in chromatin remodeling/transcription and has recently been found to be an essential and common factor that binds all three nascent arcRNAs, such as *Hsrω-n, SatIII, and Neat 1*, to form their respective nuclear bodies (43,54). Because all three arcRNAs are activated in the presence of expanded CGG repeats, it is possible that SWI/SNF complexes are brought in proximity to the chromosomal regions of the respective arcRNAs while binding to nascent arcRNAs. ISWI interacts both in vivo and in vitro with *Hsrω*-n ncRNA and stimulates its own ATPase activity via this interaction (43.). Consistent with this, we showed that expression of expanded CGG repeats results in increased binding of ISWI to Hsrω-n lncRNA, potentially influencing the cellular distribution of other associated proteins, such as Hrb87, in the formation of ω-speckles. Furthermore, we showed that ISWI repression can suppress eye phenotypes and likely structural phenotypes in polytene chromosomes in transgenic CGG repeat flies. These findings suggest that ISWI may serve as a functional intermediary between chromatin and nucleoplasmic compartments. Investigating broader transcriptional profiles in response to expanded CGG repeat expression could elucidate the connection between structural phenotypes and transcriptional regulation. In our previous studies, we observed increased expression and nuclear retention of mRNAs of stress response genes, which could reflect an initial cellular response to stress caused by expanded CGG repeats (32). Taken together, these results suggest that, at least in part, the nuclear retention of mRNAs may be related to the sequestration of certain RNA-binding proteins and/or transcription factors within *ω* speckles or other nuclear bodies. It is worth noting that in *Drosophila* expressing expanded CGG repeats, nuclear and cytoplasmic RNA foci were shown to have partial colocalization (43%) with inclusions positive for homopolymeric peptides such as ubiquitin, Hsp70, and FMRpolyG (33).

In conclusion, we present biochemical and genetic evidence to support the aberrant expression of stress-related *Hsrω-n* in CGG-mediated neurodegeneration. Furthermore, our preliminary findings showed that *SatIII* and *Neat I* were overexpressed in mammalian cells in the presence of expanded *CGG* repeats. The effects of such aberrant expression in post-mitotic neurons could include altered interactions between lncRNAs and RNA-binding proteins and chromatin remodeling factors, which could lead to changes in chromatin dynamics. It would be interesting to learn more about the genomic regions and associated mechanisms affected by expanded CGG repeats, which may provide therapeutic insights into FXTAS.

## Figure legends

**Supplementary Figure 1:**

The relative levels of CGG and Hsrω-n RNA in ten days old adult heads of indicated genotypes. *** p ≤ 0.001. n ≥ 3; Error bars indicate SEM.

**Supplementary Figure 2:**

Hrb87F protein was immunoprecipitated using P11 antibody from larval brains expressing (1) EGFP and (2) CGG_90_-EGFP. The input (10%), P11 and the IgG immunoprecipitated complexes were used for RT-PCR. Histograms show quantitative analysis of indicated transcripts obtained from P11 immunoprecipitated complexes normalized corresponding transcripts levels obtained from IgG IP complexes. *** p ≤ 0.001. n ≥ 3; Error bars indicate SEM.

## Material and Methods

### *Drosophila* strains and genetics

All flies were reared at 25°C on normal agar-cornmeal medium consisting of sugar, glucose, yeast powder, methyl paraben, and propionic acid. The wild-type strain was *w^1118^*. Transgenic flies expressing CGG_90_, CGG_60_ repeats, EGFP, and Hrb87F *(UAS-hnRNPA2/B1),* used in this study, were previously generated and described (15). Transgenic flies expressing *Hsrω (Hsrω^EP3037^, Hsrω^EP93D^)* and dsRNA against *Hsrω-n* (*Hsrω^dsRNA^*) used in this study were generously provided by Prof. Subhash Chandra Lakhotia (42)*. Elav-GAL4* (C115), *GMR-GAL4*, and *Df(3R Hrb87F/TM3, Sb^1^)* were obtained from the Bloomington Stock Center (Bloomington, Indiana, USA). A stable recombinant line expressing CGG_90_ repeats under the control of *GMR-GAL4 was* established. A stable combination line co-expressing CGG_90_ repeats and *Hsrω-n* under the control of *GMR-GAL4* was established and used for screening. Other mutant alleles used for screening were obtained from various stock centers. All crosses were grown in standard medium at 25°C.

### Tissue immunofluorescence

Tissues were dissected from larvae of the desired genotypes and processed for immunostaining, as described previously (35). The primary antibodies used to detect Hrb87F and Hsp70 proteins were mouse P11 (1:20 dilution) (67) and rat 7Fb (1:100 dilution) (68), respectively. Secondary antibodies were conjugated with Alexa Fluor 647 for anti-mouse and Alexa Fluor 546 for anti-rat antibodies (1:200; Molecular Probes, USA). Imaging was performed using a Leica SP8 Laser Scanning Confocal microscope with a 63X oil immersion objective.

### Fluorescence RNA-RNA in situ hybridization (FRISH)

SGs from late third-instar larvae were dissected in RNase-free 1× PBS and processed for RNA/RNA in situ hybridization, as previously described (69). RNA-RNA in situ hybridization was performed using a digoxigenin-labeled *Hsrω-n* riboprobe generated by in vitro transcription of the pDRM30 clone, as previously described (35). The hybridization signal was detected using a rhodamine-conjugated anti-dig antibody (1:200; Roche, Basel, Switzerland). Imaging was performed using Leica SP8 Laser Scanning Confocal microscope with 63X oil immersion objective.

### Polytene chromosome squash preparation and immunostaining

To examine the distribution of Hrb87F on the polytene chromosomes of CGG flies, SGs from actively wandering late third-instar larvae of the desired genotypes were dissected in Grace insect medium, fixed in 4% PFA for 1 min, and squashed in acetic acid and processed as previously described (70). Hrb87F on these chromosomes was detected using mouse P11 antibody (1:20 dilution; (67) and Alexa Fluor 647 anti-mouse (1:200 dilution Molecular Probes, USA) secondary antibody. Imaging was performed using Leica SP8 Laser Scanning Confocal microscope with 63X oil immersion objective.

### Scanning Electron Microscopy (SEM)

More than 50 flies were evaluated for the suppression or enhancement of the rough-eye phenotype using a dissection microscope. To confirm the phenotypes, SEM was performed on 6-8 eyes of the experimental animals with those of the controls. For SEM images, gradient concentration ethanol (25, 50, 75, and 100%) treatment was carried out to dehydrate whole flies, followed by drying with hexamethyl disilazane (Sigma). Images were analyzed using a Hitachi S-3000H scanning electron microscope.

### Viability assay in *Drosophila*

To test viability, flies of the indicated genotypes were collected and kept at 25°C for 8–10 h after eclosion. Every 2–3 days, viable flies were moved to new food vials. Dead flies were counted daily. At least three replicates were performed for each genotype.

### RNA isolation and quantitative RT-PCR

Under anesthesia, flies of the required genotypes were decapitated, and their heads were flash-frozen on dry ice. Total RNA was isolated by crushing the heads with plastic Kontes in TRIzol reagent according to the manufacturer’s protocol (Invitrogen). RNA pellets were resuspended in nuclease-free water for 10 min at 55°C and quantified using a Nanodrop (Thermo Scientific). First-strand cDNA was synthesized using Random hexamers or oligo(dT) primers for mRNA and SuperScript III (Invitrogen). The cDNA was subjected to RT-qPCR using the following gene-specific primers:

*Hsrω-n* transcript: (forward 5’-CGAAAAGGCTTATCCTCTTGGTAAA-3’ and reverse 5’-AAGGATAATGATTAAGGTAATCGGG-3’)(71).
*18s* transcript: (forward:5’-CGGCTACCACATCCAAGGAA-3’ and reverse: 5’-GCTGGAATTACCGCGGCT -3’).
*EGFP* transcript: (forward 5’-CTGCTGCCCGACAACCA-3’ and reverse: 5’-GAACTCCAGCAGGACCATGTG-3).’
*Sat III*: (forward 5’-TATGAATTCAATCAACCCGAGTGCAATCGAA-3’ and reverse:: 5’-TATGGATCCTTCCATTCCATTCCTGTACTCG-3’)
*Neat 1_1 (Neat 1S):* (forward: 5’-GTGGCTGTTGGAGTCGGTAT-3’and reverse: 5’ TAACAAACCACGGTCCATGA -3’)
*β-Actin*: (forward: 5’-TGAGAGGGAAATCGTGCGTG-3’ and reverse: 5’TGCTTGCTGATCCACATCTGC-3’
*hSNF2h*: (forward: 5’ TAC AAA CAA CTG CCT TGG G 3’; and reverse: 5’ TTG GAG GCA AAC TCT TTT CAA -3’)
*Hsp70:* (forward: 5’-CTCGTCGGCGGATCCA-3’ ; and reverse: 5’GGAAGAACTCCTGCAGCAGACT-3’)
Hrb87F: (forward primer 5’-CACGTACTCCCAGTCGTACAT-3’; and reverse: 5’GCAGGCACTCTTCATCGTGA-3’
ISWI: (forward primer 5’-AAGAGTCCCACGAAGCCTAAG-3’; and reverse: 5’-GTGAGGCATCGAAGCGAAAGA-3’
Rox1: (forward primer 5’-GGATTTCCATGTTTGTTGCTGC-3’; and reverse: 5’ATACGCCTGAGCCGATTCAC-3’
U4: (forward primer 5’-CCTGAGGTGCGGTTATTGCT-3’; and reverse: 5’CCCTCTCGGGCTTCCAAAAA-3’
hSNF2h: (forward primer 5’-TGCAGGTTGGATGGTCAGACAC-3’; and reverse: 5’GTCGCAAGATTGATGCCAAGACC-3’

The 7500 Standard Real-Time PCR System was used to run real-time PCR in triplicate using gene-specific primers and SYBR Green PCR Master Mix (Applied Biosystems). For fly samples, 18s was used as an endogenous control. β-actin was used as an endogenous control for *SatIII* and *Neat1*. Relative transcript levels were computed using the 2^−^ΔΔCt^ method, and the control samples were calibrated.

### Transcriptomic sequencing and data analysis

Cerebellar tissues of FXTAS mice and littermate controls were collected for RNA extraction by using TRIzol® reagent. A total of 1μg RNA per sample was processed for library preparation with the NEBNext® Ultra™ II Directional RNA Library Prep Kit with NEBNext® rRNA Depletion Kit (RiboZero Plus kit). Libraries were sequenced on an Illumina HiSeq platform by targeting 100M reads per sample. RNASeq reads were mapped to the mouse genome (mm10) by TopHat2 with default parameters, followed by gene expression quantification using FeatureCounts. Differential gene expression analysis was performed using DESeq2 R library.

### Cross-Linking RNA Immuno-Precipitation (CLIP-RIP)

CLIP-RIP was performed as previously described (67), with some modifications. Two hundred micrograms of snap-frozen larval brains of desired genotypes or 100 µg of volume packed S2 cells (3 days post-transfection) were crushed with Kontes, resuspended, and fixed in DEPC-treated 1X PBS containing 1% formaldehyde for 20 min under rotation. Cross-linking was stopped by the addition of glycine (pH 7.0) to a final concentration of 0.25 M for 5 min at room temperature followed by centrifugation at 1100× g for 2 min at 4 °C to collect the pellet. The pellet was washed with cold DEPC-treated 1× PBS, resuspended in 10 ml final volume of nuclear extraction buffer (NEB), and centrifuged at 3220× g for 5 min at 4 °C to collect the cross-linked pellet. The pellet was subsequently resuspended in 2 mL RIPA buffer. The cross-linked pellet prepared from the larval brains underwent an additional step of mechanical shearing by pipetting in and out and flowing through a 25GX 5/8-inch needle until the lysate easily passed through the needle. Finally, solubilization was achieved by sonication, as previously described (45), and the debris was pelleted at 16,000 × g for 5 min at 4°C. The supernatant was collected to calculate the protein concentration using the Bradford assay. An aliquot of solubilized cell lysate (250 μL)) was mixed with Protein A-magnetic beads (25μL) along with non-specific competitor tRNA (final concentration of 100ug/mL) under rotation for 1 h at 4°C, followed by centrifugation to remove the beads. The precleared supernatant was used for subsequent processes. Protein A-magnetic beads (25μL) were resuspended in 1XPBS and incubated with 2μg of anti-Hrb87F (P11) or generic mouse IgG (Cell Signaling Technology) overnight at 4°C with gentle agitation. The beads were then centrifuged for 2 min at 4000 × g, washed three times with 500μl 1XPBS and finally incubated with 200μg of total protein lysate in 125μl final volume of IP buffer overnight at 4°C, followed by several brief washes with 1.2mL of IP buffer for 10min at room temperature. Finally, the bound material was eluted with 30μl of TEL buffer treated with 600μL of TRIzol (Invitrogen, Thermo Fisher Scientific) for RNA extraction as per the manufacturer’s recommendations. Buffers: Nuclear extraction buffer (NEB) (15 mM Hepes pH = 7.0, 5mM MgCl_2_, 0.2 mM EDTA, 0.5 mM EGTA, 10mM KCl, freshly added 350mM sucrose and 1mM DTT and 0.1%Tween-20), RIPA buffer (50mM Tris-HCl pH 7.5, 1% NP40, 0.5% Na-deoxycholate, 0.05% SDS, 1 mM EDTA, 150mM NaCl and protease inhibitors (complete, mini, EDTA-free protease inhibitor cocktail tablet, Roche), IP buffer: (10mM Hepes-KOH, pH 8.0, 100mM NaCl, 10% glycerol, 0.05% Tween 20, 100μg/ml PMSF, complete protease inhibitors-Roche 1X). TEL buffer 200mM Tris-HCl, pH 7.4, 25mM EDTA, pH 8.0, 100mM LiCl, 1% SDS).

### DNA constructs

Mammalian expression plasmids pcDNA3.1, expressing 20, 40, 60, or 99 CGG repeats, have been described previously (13). 99 CGG repeats were subcloned from pcDNA3.1-CGG99X into the metallothionein-inducible Drosophila expression pRMHa3-GFP. Similarly, the conjugated catalytic domain of ISWI was subcloned into pRMHa3-GFP.

### Cell culture, transfection, and treatments

Human embryonic kidney 293T (HEK293T) cells were grown on poly-L-lysine-coated 6-cm dishes in DMEM (Hyclone) supplemented with 10%FBS, 0.5 mM L-glutamine, and 100 IU/mL penicillin streptomycin at 37°C with 5% CO_2_ and transfected on day 3. Drosophila Schneider 2 (S2) were cultured in Schneider’s Drosophila Medium (Gibco BRL/Invitrogen) supplemented with 10% fetal bovine serum and gentamicin/penicillin at 25°C and transfected 24 h later. Transfection was done by using Effectene (Qiagen) or PEI (G-biosciences) transfection reagent according to the manufacturer’s protocol

### SDS-PAGE and Western blot analysis

Protein lysates treated with equal volume of SDS-PAGE 2x Laemmeli buffer were electrophoretically separated on a 12% SDS–PAGE gel and subsequently transferred to a polyvinylidene fluoride membrane (Millipore, Bedford, MA, USA). Western blotting was performed using Tris buffer (pH 8.3). The following antibody dilutions were used: mouse anti-β-tubulin (E7, Developmental Studies Hybridoma Bank; 1:200), rabbit anti-GFP (Cell Signaling Technology, #2956; 1:1000), and anti-Hrb87F (P11; 1:100) developed with species-specific IRDye-conjugated secondary antibodies (LI-COR Biosciences; 1:5000, and analyzed using an Odyssey Infrared Imaging System (LI-COR Biosciences). Relative quantification of the desired protein was estimated using the ratio of the band intensities of the target proteins to that of the reference (β-tubulin). Densitometry measurements were performed in ImageJ using scanned films with the same exposure times across multiple experiments.

### Statistical method

The log-rank (Mantel-Cox test) tests were used to compare the differences between the survival curves using the GraphPad Prism software. To determine the significance and p-values, statistical analysis was performed using one-way analysis of variance (ANOVA) with Tukey’s post-hoc tests and Bonferroni correction for multiple comparisons (GraphPad Prism software). The unpaired t-test was used for two-sample comparisons. All data are presented as the mean ± standard error of the mean (mean ± SEM).

## Acknowledgments

The authors thank Dr. Nicolas Charlet Berguerand (Institute of Genetics, Molecular and Cellular Biology IGBMC, France) for providing useful reagents for this study. We especially thank Prof. Harald Saumweber (Institute of Biology, Humboldt University of Berlin, Germany) and Prof. Subhash Chandra Lakhotia (Banaras Hindu University, Varanasi, India) for providing the anti-P11 antibody and suggestions in designing the confocal microscopy experiments. The transgenic *Drosophila* lines used in this study, including Hsrω^EP3037^, Hsrω^EP93D^, and Hsrω^dsRNA^, were generously provided by Prof. Subhash Chandra Lakhotia. The authors would like to express their gratitude to the USIC facility at the University of Kashmir for their technical assistance with SEM. Thanks to SATHI-CDC (Banaras Hindu University, Varanasi, India) for the confocal facility. The authors would also like to thank the Qurashi laboratory members for their help and critical reading of this work.

## Author Contributions

A.Q. conceived and designed the experiments. N. A., M.M.R., and A.Q. performed experiments. A.Q., M.M.R., N.A., and P.J. analyzed the data. A. Q. wrote the manuscript.

