## Supplementary Figures for "Aberrant expression of stress-related Hsrω-n lncRNA contributes to CGG repeat-mediated toxicity in a *Drosophila* model of FXTAS"

S1

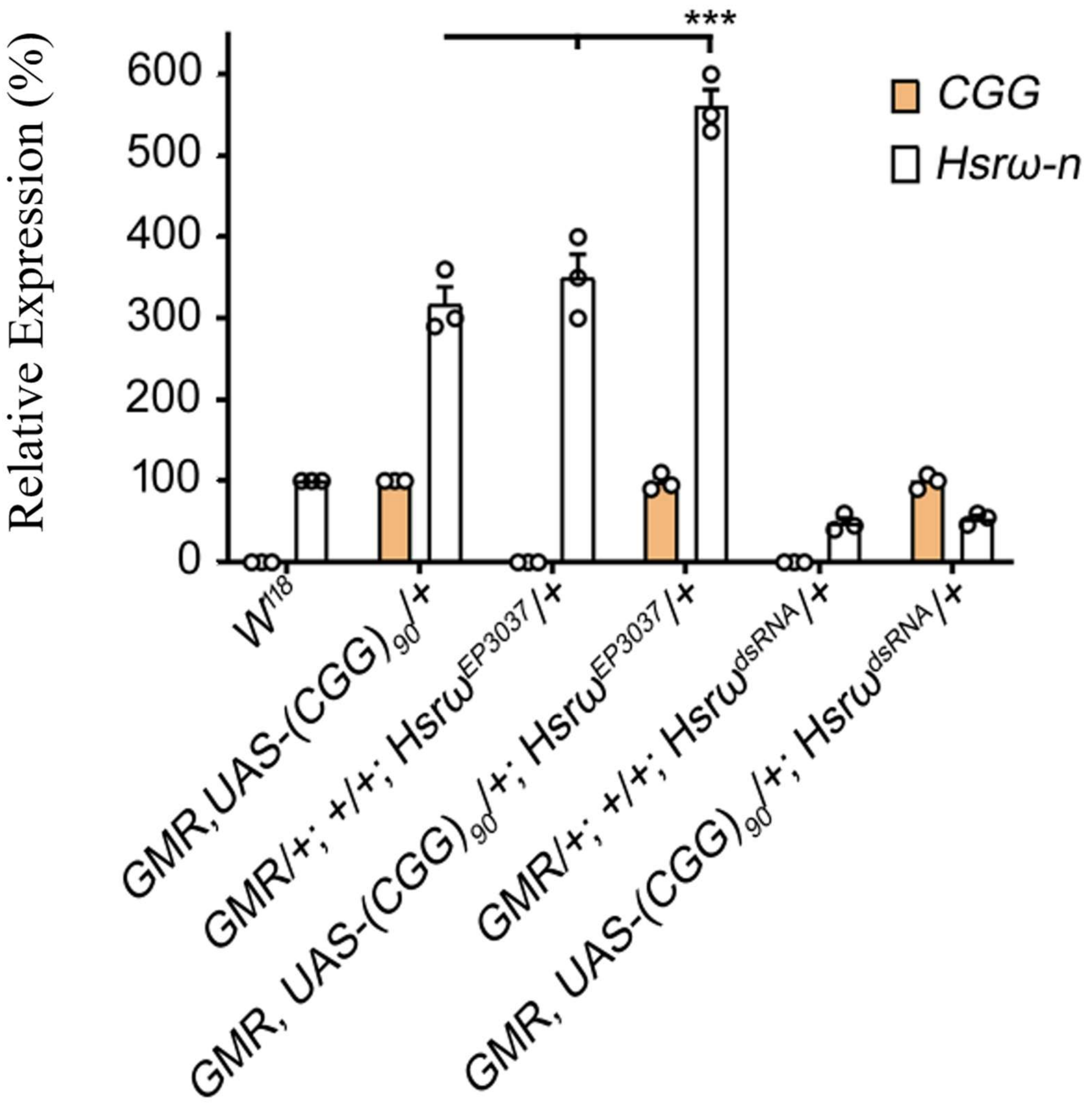


**Supplementary Figure 1:**

The relative levels of CGG and Hsrω-n RNA in ten days old adult heads of indicated genotypes. *** $p \leq 0.001$. $n \geq 3$; Error bars indicate SEM

S2

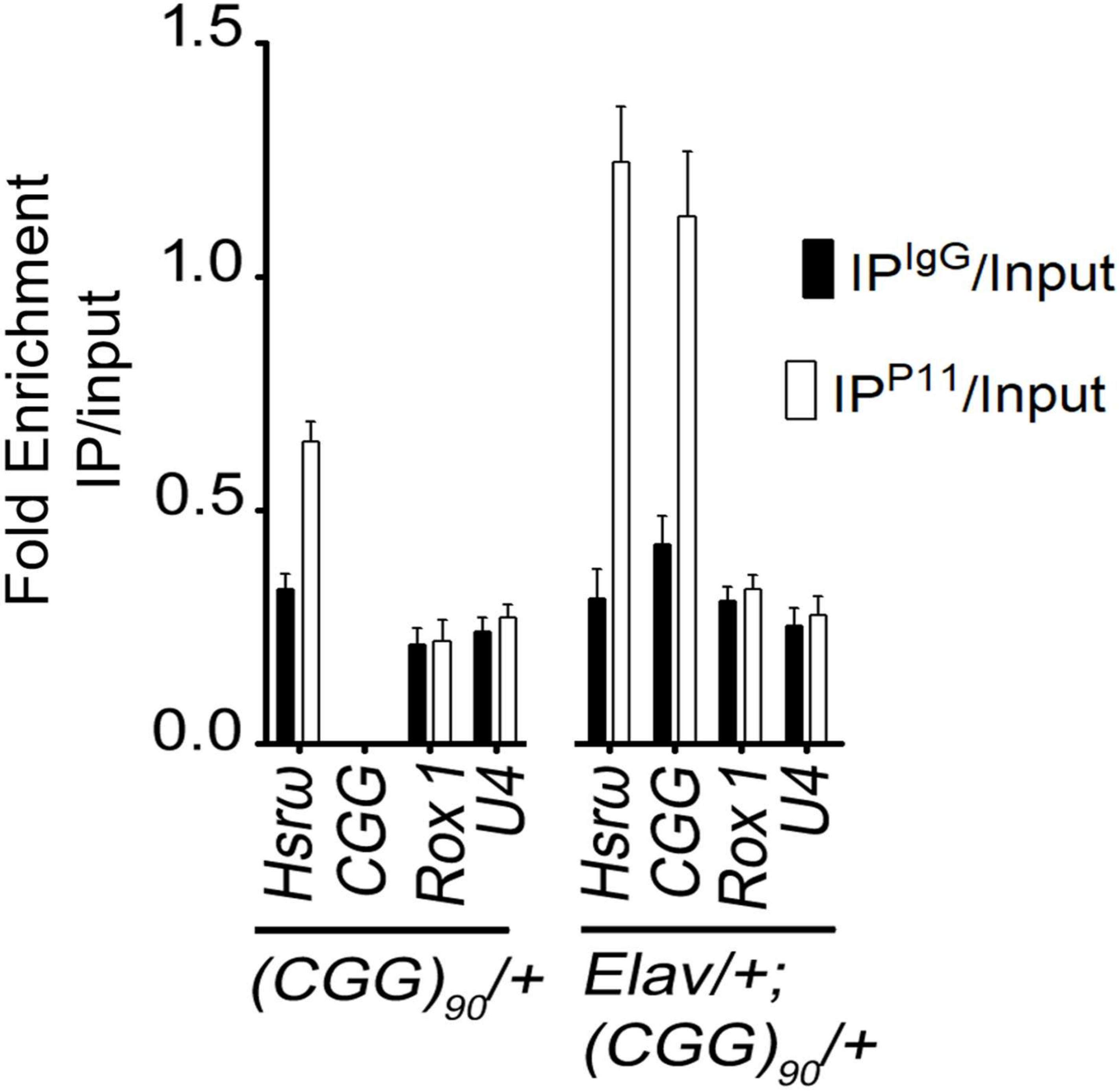


**Supplementary Figure 2:**

Hrb87F protein was immunoprecipitated using P11 antibody from larval brains expressing (1) EGFP and (2) $CGG_{90}$-EGFP. The input (10%), P11 and the IgG immunoprecipitated complexes were used for RT-PCR. Histograms show quantitative analysis of indicated transcripts obtained from P11 immunoprecipitated complexes normalized corresponding transcripts levels obtained from IgG IP complexes. *** $p \leq 0.001$. $n \geq 3$; Error bars indicate SEM.
